# Read&Tag enables antibody-independent combinatorial and multifactorial chromatin profiling

**DOI:** 10.64898/2026.11.19.763725

**Authors:** Marius Schneider, Lara C. Kopp, Christa Schwarzlose, Namisha R. Bohara, Chondamma Bollachettira, Aline Campos Sparr, Lukas A. Huschet, Maria S. Robles, Rodrigo Villaseñor

## Abstract

Genome-wide chromatin profiling methods based on in situ tagmentation have expanded access to epigenomic information but generally rely on antibodies, thereby limiting their multifactorial capacity and the direct measurement of combinatorial chromatin states. Here, we introduce Read&Tag, in which engineered chromatin reader domains or intrabodies are fused directly to the Tn5 transposase to target specific chromatin features in situ. Read&Tag profiles active and repressive histone modifications, DNA methylation, and elongating RNA polymerase II, with signal amplitude increasing with target abundance and remaining robust in frozen tissue and low-input samples. Because the targeting module is a protein of defined structure and affinity, two specificities can be combined within a single complex. Tagmentation by such a dual-specificity reader requires H3K4me3 and H3K27me3 to be present together in the same chromatin fragment. The co-occurrence of both marks thus becomes a precondition of the measurement rather than an inference drawn from independently measured profiles. Combining barcoded reader-Tn5 complexes with primer-timed signal augmentation enables the measurement of four chromatin features in a single reaction. Read&Tag therefore provides a modular, antibody-independent platform for combinatorial and multifactorial profiling.

## Introduction

Chromatin profiling methods have become essential techniques for understanding gene regulation, cell identity, and disease by mapping the genomic distribution of histone modifications, transcription factors, chromatin-associated proteins, and chromatin accessibility. In the past decade, significant advances have simplified and scaled epigenomic assays ^1^, including the development of in situ approaches such as CUT&RUN ^2^, ChIL-seq ^3^, CUT&Tag ^4^, ACT-seq ^5^, and CoBATCH ^6^. Subsequent transposase-based methods have extended this principle to multiplexed profiling of several targets in one reaction using barcoded antibody-Tn5 complexes ^7–9^, direct antibody-Tn5 conjugation ^10^, replacement of the protein A-antibody tether with recombinant binders, either nanobodies raised against the primary antibody ^11,12^, or binders directed at a genetically encoded tag on the target ^13,14^. In all of these approaches, target recognition is ultimately performed by an antibody or by an epitope introduced into the genome.

These methods have significantly reduced input requirements and improved signal-to-noise compared with traditional chromatin immunoprecipitation followed by sequencing (ChIP-seq) ^15^, thereby enabling chromatin profiling with limited sample amounts. Despite these advances, most current chromatin profiling techniques still primarily rely on antibodies, which are limited by availability, specificity, and reproducibility for chromatin-associated targets. Antibody performance can differ widely across epitopes, experimental conditions, and sample types, often requiring extensive optimization ^16,17^. Additionally, mapping combinatorial chromatin states is not easily achievable with standard antibody profiling. Lastly, transposase-based methods are inherently sensitive to chromatin accessibility ^18^, which may complicate the interpretation of chromatin states. These limitations motivate the development of alternative targeting strategies that decouple chromatin recognition from antibody reagents while preserving the efficiency and flexibility of in situ tagmentation.

In cells, chromatin modifications are recognized by reader domains that bind specific histone marks or DNA modifications with defined specificity and affinity ^19^. Because these domains can be produced recombinantly and rationally modified ^20–28^, they provide a strong alternative to antibodies as targeting reagents for chromatin profiling. However, a general strategy for coupling chromatin readers to transposase-mediated library preparation in a modular and extensible manner has been lacking.

Here, we introduce Read&Tag, a chromatin profiling strategy based on reader-guided transposition, in which engineered chromatin readers (eCRs) ^26,27^ are fused directly to the Tn5 transposase to mediate in situ insertion of DNA sequencing adapters at specific chromatin features. By replacing antibody-mediated recruitment with recombinant eCR-Tn5 fusions, Read&Tag offers a modular, extensible platform for chromatin profiling that is independent of antibodies and amenable to systematic combinatorial and multifactorial profiling. This design allows direct assessment of reader-chromatin specificity and rational optimization of targeting performance through domain engineering. We demonstrate that Read&Tag enables specific profiling of diverse chromatin features with a signal amplitude that scales with target abundance, including active and repressive histone modifications, DNA methylation and transcription-associated RNA polymerase II phosphorylation. Because the targeting module is a reader domain of defined structure and affinity, two specificities can be combined within a single complex, so that tagmentation requires two histone marks to be present together in the same chromatin fragment, here H3K4me3 and H3K27me3. Co-occurrence of both marks thereby becomes a precondition of the measurement rather than an inference drawn from independently measured profiles. Through comparative analyses of dual-domain reader architectures and targeted reader mutations, we show that genomic localization is driven by bona fide reader-chromatin interactions. Read&Tag recapitulates canonical chromatin landscapes across multiple contexts, including frozen tissue and low-input samples, while preserving high specificity. The same modularity extends to multiplexing. By combining barcoded reader-Tn5 complexes with primer-timed signal augmentation, Multi-Read&Tag enables balanced, multifactorial profiling of four distinct chromatin features in a single reaction, with no loss of specificity and no signal transfer between channels above the detection limits of the assay. Our results position Read&Tag as an antibody-independent alternative to current chromatin profiling methods that retains the workflow and sensitivity of transposase-based library preparation while replacing the antibody and protein A-Tn5 module with a single recombinant protein.

## Results

### Design and characterization of eCR-Tn5 transposase fusions for epigenomic profiling

To enable selective epigenomic profiling in mammalian cells, we developed a strategy based on recombinant engineered chromatin readers fused to a Tn5 transposase. We designed recombinant eCR-Tn5 fusion proteins incorporating either native ^26^ or engineered reader domains ^27^ that recognize defined chromatin features, including H3K4me3, H3K27me3, H3K9me3, and 5-methylcytosine (5mC) (Fig. 1a). To systematically assess the impact of reader valency, we generated both single-domain (1x) and dual-domain (2x) eCR-Tn5 constructs for each chromatin target (Fig. 1b).

**Figure 1.**
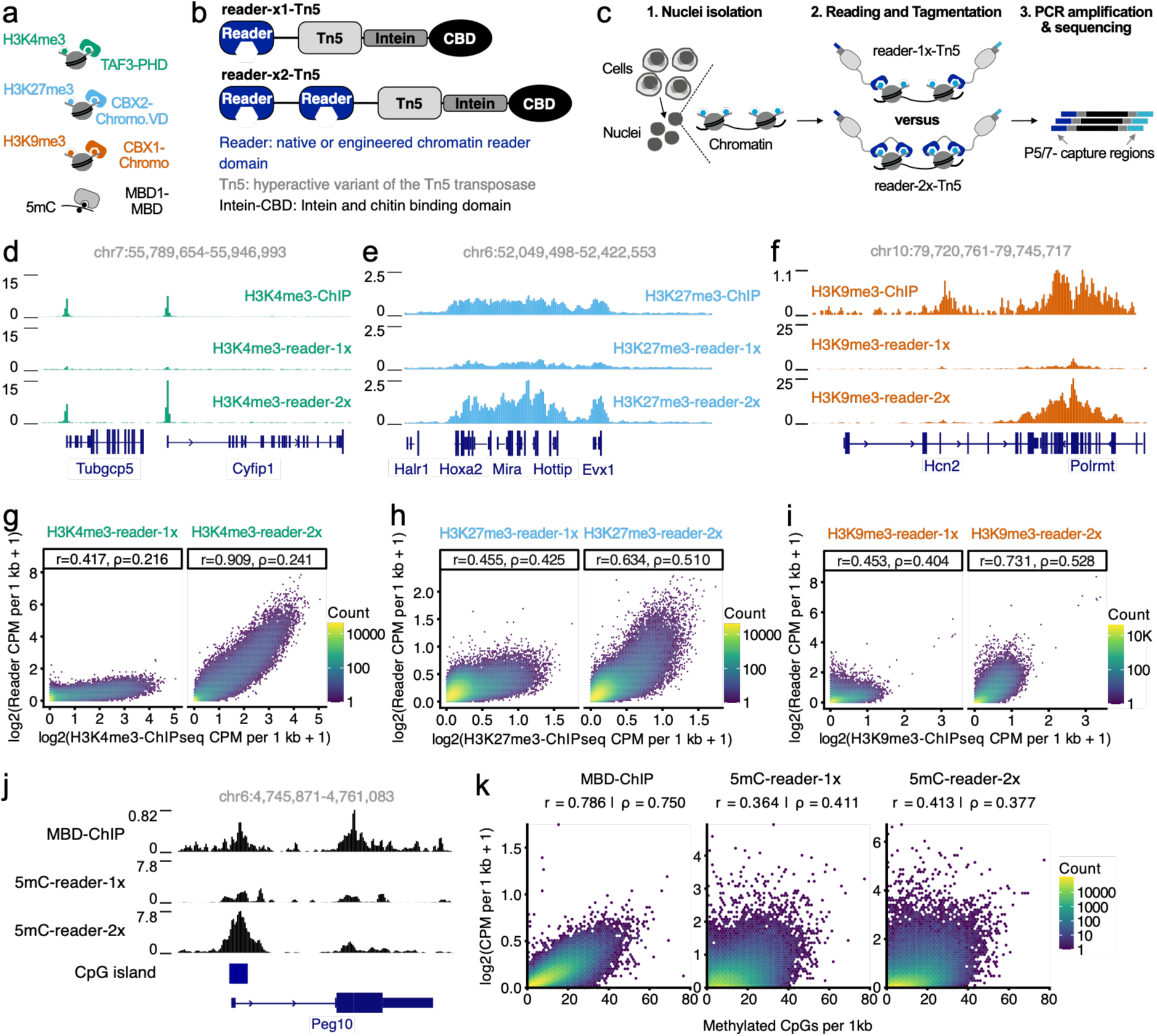
Design and characterization of eCR-Tn5 transposase fusions for epigenomic profiling. Chromatin domains used in Read&Tag, experimental workflow, and representative comparisons between eCR-Tn5 profiles and ChIP-seq datasets. **(a)** Schematic overview of native or engineered chromatin reader domains used in this study and their cognate chromatin targets, including the H3K4me3-specific PHD domain of mouse TAF3, an engineered H3K27me3-specific chromodomain (CBX2.VD variant), the H3K9me3-specific chromodomain of mouse CBX1/HP1beta, and the methyl-CpG-binding domain (MBD) of murine MBD1. **(b)** Schematic representation of single-domain (reader-x1-Tn5) and dual-domain (reader-x2-Tn5) fusion protein architectures. Chromatin reader domains are fused to a hyperactive Tn5 transposase, followed by a C-terminal intein-chitin-binding domain (CBD) used for affinity purification. **(c)** Outline of the Read&Tag workflow. Nuclei are isolated and immobilized on Concanavalin A-coated beads, followed by incubation with single-domain or dual-domain reader-Tn5 fusion proteins. Reader-guided tagmentation integrates sequencing adapters at targeted chromatin features, after which libraries are PCR-amplified and sequenced. **(d, e, f, j)** Genome browser tracks showing the mean signal (reader-x1-Tn5, n=2; reader-x2-Tn5, n=3) at representative loci for H3K4me3 (d), H3K27me3 (e), H3K9me3 (f), and 5-methylcytosine DNA methylation (j), comparing conventional ChIP-or MBD-ChIP-seq profiles with Read&Tag profiles generated using single-domain (x1) and dual-domain (x2) reader-Tn5 fusions. **(g, h, i)** Genome-wide comparison of mean Read&Tag signal (reader-x1-Tn5, n=2; reader-x2-Tn5, n=3) to corresponding reference datasets (n=2) for each chromatin feature. Scatter plots show 1-kb binned log2-transformed CPM values (log2(x+1)) for Read&Tag, ChIP-seq or MBD-ChIP-seq, with color indicating bin density. Panels correspond to H3K4me3 (g), H3K27me3 (h), and H3K9me3 (i) with separate plots for single-domain and dual-domain reader configurations. Pearson correlation (r) and Spearman correlation (ρ) were calculated across 2,220,883 genomic windows for each feature. **(k)** Genome-wide relationship between MBD-Tn5 signal and CpG methylation density. Scatter plots show 1-kb binned log2-transformed CPM values (log2(x+1)) versus the number of methylated CpGs per kilobase for MBD-ChIP-seq (ref. 26), 5mC-reader-1x-Tn5, and 5mC-reader-2x-Tn5 datasets. Pearson correlation (r) and Spearman correlation (ρ) were calculated across 2,323,263 genomic windows. Differences in window counts reflect feature-specific filtering (e.g., CpG coverage).

All eCR-Tn5 fusion proteins were expressed in *E. coli* and purified using an intein-mediated one-step affinity purification strategy ^29^. All constructs carried a C-terminal intein-chitin-binding domain (CBD) tag, enabling affinity purification on chitin resin (Supplementary Fig. 1a). On-column intein cleavage induced by dithiothreitol (DTT) released the eCR-Tn5 fusion proteins from the affinity tag, yielding pure and active transposase suitable for chromatin profiling (Supplementary Fig. 1b). Purified eCR-Tn5 fusions were pre-loaded with sequencing adapters and incubated with permeabilized nuclei immobilized on Concanavalin A-coated paramagnetic beads. Upon activation by magnesium ions, reader-guided tagmentation integrated sequencing adapters into genomic regions bound by the cognate chromatin reader. The tagmented genomic DNA was subsequently extracted and amplified by PCR to generate indexed sequencing libraries, which were pooled and subjected to paired-end Illumina sequencing (Fig. 1c).

We first evaluated the performance of single-domain and dual-domain eCR-Tn5 proteins by profiling H3K4me3, H3K27me3, H3K9me3, and 5mC in 500,000 mouse embryonic stem cells (mESCs). Genome browser tracks at representative loci showed that dual-domain eCR-Tn5 fusions more closely recapitulated reference ChIP-or methyl-CpG-binding-domain-ChIP-seq (MBD-ChIP-seq) ^26^ profiles than their single-domain counterparts for all chromatin marks tested (Fig. 1d-f, j). This improvement was evident for both active chromatin (H3K4me3) and repressive chromatin states (H3K27me3 and H3K9me3), as well as for DNA methylation (5mC). To quantify these observations genome-wide, we computed Pearson (r) and Spearman (ρ) correlations between eCR-Tn5 signal and matched reference datasets across 2,220,883 non-overlapping 1-kb autosomal windows. Dual-domain (2x) constructs increased genome-wide Pearson correlations by 0.18-0.49 relative to single-domain (1x) designs (Fig. 1g-i), while retaining comparable fragment length distributions across constructs (Supplementary Fig. 1c-d). The genome-wide Pearson improvement was largest for sharp, narrowly distributed marks such as H3K4me3 (Fig. 1g), where the signal-bearing fraction of the genome is small and quantitative agreement at peaks dominates the metric. Because rank-based Spearman correlation across all 1-kb windows is dominated by the large background fraction in which neither construct carries informative signal, ρ improvements at the genome scale are more modest. Restricting the analysis to windows overlapping reference consensus peaks for these marks, where biologically meaningful signal is concentrated, both Pearson and Spearman correlations increased substantially with the 2x design (Supplementary Fig. 1e; Supplementary Table S1).

For 5-methylcytosine, the 5mC-reader-2x binds individual methylated CpG dinucleotides with affinity that scales with local mCpG density. This is consistent with the low-resolution and semi-quantitative readout of DNA methylation by MBD-ChIP-seq (Fig. 1k). Next, we compared MBD-Tn5 against the orthogonal WGBS gold standard. After matching all MBD-Tn5 libraries to a common sequencing depth to exclude depth as a confounder, valency-driven improvement increased specifically at high methylation density (16-25 methylated CpGs/kb; Δr = 0.047, cross-replicate range [0.004, 0.088]) and was essentially unchanged from the unmatched analysis, whereas the constructs were indistinguishable at lower densities (Supplementary Fig. 1f). Notably, the correlation between the single-domain reader and WGBS density did not exceed mean r = 0.25 at the most densely methylated regions (26+ methylated CpGs/kb), while the dual-domain construct reached r = 0.33, showing that doubled valency is necessary for semi-quantitative methylation readout through Tn5 transposition at high methylation density (Supplementary Fig. 1g).

Together, and in agreement with previous studies ^22–24,26^, these results indicate that increasing reader valency improves signal concordance with reference data both globally and at biologically meaningful loci, and, for 5-methylcytosine specifically, this improvement is density-dependent, consistent with avidity-enhanced cooperative binding to methylated CpGs. Hereafter, Read&Tag mainly refers to eCR-guided transposition using dual-domain constructs.

### Specificity of dual-domain eCR-Tn5 fusions depends on intact reader-chromatin interactions

To directly test whether chromatin targeting by dual-domain eCR-Tn5 fusions depends on specific reader-histone interactions, we introduced amino-acid substitutions previously shown or predicted to disrupt binding of the respective reader domains: TAF3-D890A/W891A ^30^, CBX1-W42A/F45A ^31^, and CBX2-W33A/W36A. Genome browser tracks at representative loci revealed that these mutations abolished enrichment of Read&Tag (RnT) signal at regions marked by the corresponding histone modifications, including H3K4me3, H3K27me3, and H3K9me3 (Fig. 2a-c). Consistent with these qualitative observations, genome-wide correlation analysis showed that mutant Read&Tag profiles exhibited a substantial reduction of correlation with matched ChIP-seq or CUT&Tag datasets (Fig. 2d), indicating loss of eCR-guided transposition.

**Figure 2.**
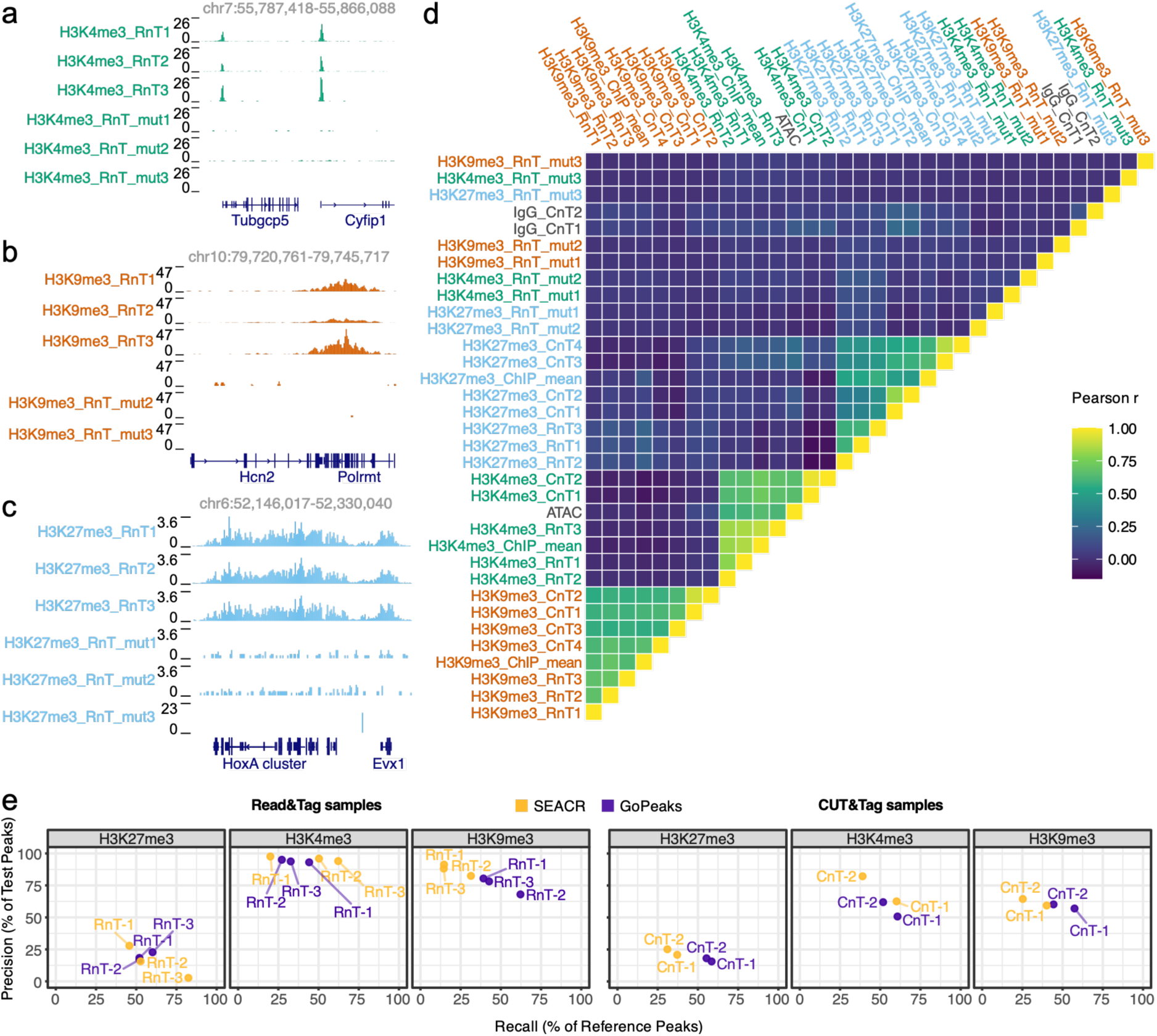
Reader-dependent specificity of Read&Tag revealed by mutational controls and genome-wide benchmarking. Assessment of Read&Tag specificity using targeted reader mutations. **(a-c)** Genome browser tracks at representative loci showing profiles for H3K4me3 (a), H3K27me3 (b), and H3K9me3 (c). For each histone modification, Read&Tag (RnT) profiles are shown, generated using the corresponding dual-domain eCR-Tn5 fusion in either wild-type or reader-mutant form. Gene annotations are shown below each locus, and genomic coordinates are indicated above. **(d)** Pairwise Pearson correlation heatmap comparing genome-wide signal distributions across Read&Tag biological replicates (n=3) and reference ChIP-seq (mean, n=2) and CUT&Tag (CnT, n=2-4) datasets (refs. 7, 32-34) for the three histone modifications. Each cell represents the Pearson correlation coefficient between two samples, with color indicating correlation strength. **(e)** Precision-recall plots summarizing peak recovery for Read&Tag and CUT&Tag (ref. 33) datasets across the three ChIP-seq-defined histone modification reference peak sets. Points represent individual datasets, with axes showing recall (percentage of reference peaks recovered) and precision (percentage of test peaks overlapping reference peaks). Data are displayed separately for Read&Tag and CUT&Tag samples, and points are labeled by biological replicate and peak-calling strategy (GoPeaks or SEACR).

We next quantified enrichment using the fraction of reads in peaks (FRiP). Mutant constructs showed a pronounced reduction in FRiP across all histone marks, approaching background levels observed for input controls (Supplementary Fig. 2a). In contrast, wild-type dual-domain eCR-Tn5 fusions displayed strong enrichment relative to matched ChIP-seq or CUT&Tag ^7,32–34^ datasets. Notably, for H3K27me3 and H3K9me3, enrichment remained low in accessible chromatin regions when considering non-duplicated reads, indicating that signal is not driven by chromatin accessibility but by specific reader-histone interactions (Supplementary Fig. 2a). H3K9me3 Read&Tag samples additionally showed modest sensitivity to PCR duplication in the FRiP analysis, consistent with signal arising from highly enriched, low-complexity genomic regions. While two published CUT&Tag datasets targeting H3K27me3 ^7,32^ exhibited higher FRiP values than Read&Tag in this comparison, these profiles also showed increased signal in accessible chromatin, consistent with a contribution from accessibility-driven activity under the reported experimental conditions (Supplementary Fig. 2a). Together, these results demonstrate that the specificity and genomic localization of dual-domain eCR-Tn5 fusions strictly require intact reader-chromatin interactions and that the observed signal predominantly reflects eCR-guided, rather than nonspecific accessibility-driven transposition.

To further evaluate specificity independently of enrichment magnitude, we performed a precision-recall analysis using MACS2-defined ^35^ ChIP-seq peaks as reference sets. Test peaks from Read&Tag and CUT&Tag datasets were called using SEACR ^36^ and GoPeaks ^37^, with the corresponding mutant-reader or IgG samples used as background controls. Precision-recall benchmarking revealed method-dependent differences across histone marks (Fig. 2e). SEACR yielded higher precision but lower recall, consistent with a conservative peak-detection approach that captures only the strongest signal regions. In contrast, GoPeaks provided a more balanced performance across all histone marks, achieving intermediate-to-high precision and recall, with peak numbers and widths consistent with expected chromatin domain architectures (Supplementary Fig. 2b-c). Together, these results demonstrate that dual-domain eCR-Tn5 fusions achieve highly specific chromatin targeting that depends on intact reader-histone interactions and predominantly reflects targeted reader-guided transposition.

To test whether Read&Tag signal amplitude scales with target abundance, we applied it in PRC1-degron cells, in which H3K27me3 erodes gradually after RING1B degradation ^38^. We added *Drosophila melanogaster* nuclei as an exogenous chromatin spike-in and performed Read&Tag after 0, 8 and 24 hours of auxin treatment. Loss of the PRC1 catalytic product H2AK119ub1 confirmed degradation (Supplementary Fig. 2d). These data and the spike-in-calibrated H3K27me3 ChIP-seq of Dobrinić et al.^38^ from the same cells were processed through an identical pipeline and analyzed both with and without spike-in normalization, across 2,745 H3K27me3 domains defined from the reference ChIP-seq independently of Read&Tag. Both assays report loss at 86-100% of domains depending on timepoint and normalization, within the range of replicate-to-replicate agreement in either assay. CPM-and spike-in-normalized Read&Tag describe the same trajectory of H3K27me3 loss, and both track the calibrated ChIP-seq series to within 0.8-1.1-fold at both timepoints (Supplementary Fig. 2e)

### Read&Tag reproduces canonical chromatin modification patterns in human cells

Having established that chromatin targeting by Read&Tag depends on specific reader-histone interactions, we next assessed its performance in cultured human cells. We profiled three histone modifications and 5mC in 250,000 nuclei from the breast cancer cell line MCF7. Genome browser inspection revealed robust, mark-specific enrichment: sharp promoter-associated peaks for H3K4me3, broad facultative and constitutive heterochromatin domains for H3K27me3 and H3K9me3, respectively, and focal enrichment at hypermethylated loci for 5mC (Supplementary Fig. 3a-d). Genome-wide correlation analysis showed strong concordance between biological replicates and clear clustering by modification. As in mouse ES cells, mutant Read&Tag profiles showed markedly reduced correlation with matched ChIP-seq ^39^ and CUT&Tag ^40^ datasets, with residual signal redistributing to accessible chromatin (Supplementary Fig. 3e,f), consistent with loss of eCR-guided transposition. Finally, FRiP analysis showed mark-appropriate enrichment within reference peak sets across all marks, high for focal H3K4me3 and lower for the broad H3K27me3 and H3K9me3 domains, at levels on par with published CUT&Tag ^40^ and MeDIP ^41^ samples, with little sensitivity to PCR duplication or accessibility-driven transposition (Supplementary Fig. 3f). Read&Tag therefore recapitulates the human chromatin modification landscape at a quality comparable to ChIP-seq or CUT&Tag for histone marks and MeDIP for 5mC.

### Read&Tag faithfully captures canonical histone modification patterns in frozen liver tissue

To assess the versatility of Read&Tag, we profiled histone modifications in 500,000 nuclei enriched from mouse liver tissue ^42^ stored at −80 °C for over two years. Genome browser inspection revealed robust and mark-specific enrichment patterns for H3K4me3, H3K27me3, and H3K9me3, corresponding to sharp promoter-associated peaks, broad facultative heterochromatin, and broad constitutive heterochromatin domains, respectively (Fig. 3a-c). These patterns closely matched published ChIP-seq profiles from flash-frozen liver tissue ^43,44^, indicating that Read&Tag preserves histone modification specificity in frozen tissue samples. Consistent with these observations, pairwise correlation analysis showed strong concordance between biological replicates and clear clustering by histone modification rather than by sample identity, with minimal cross-mark correlation (Fig. 3d).

**Figure 3.**
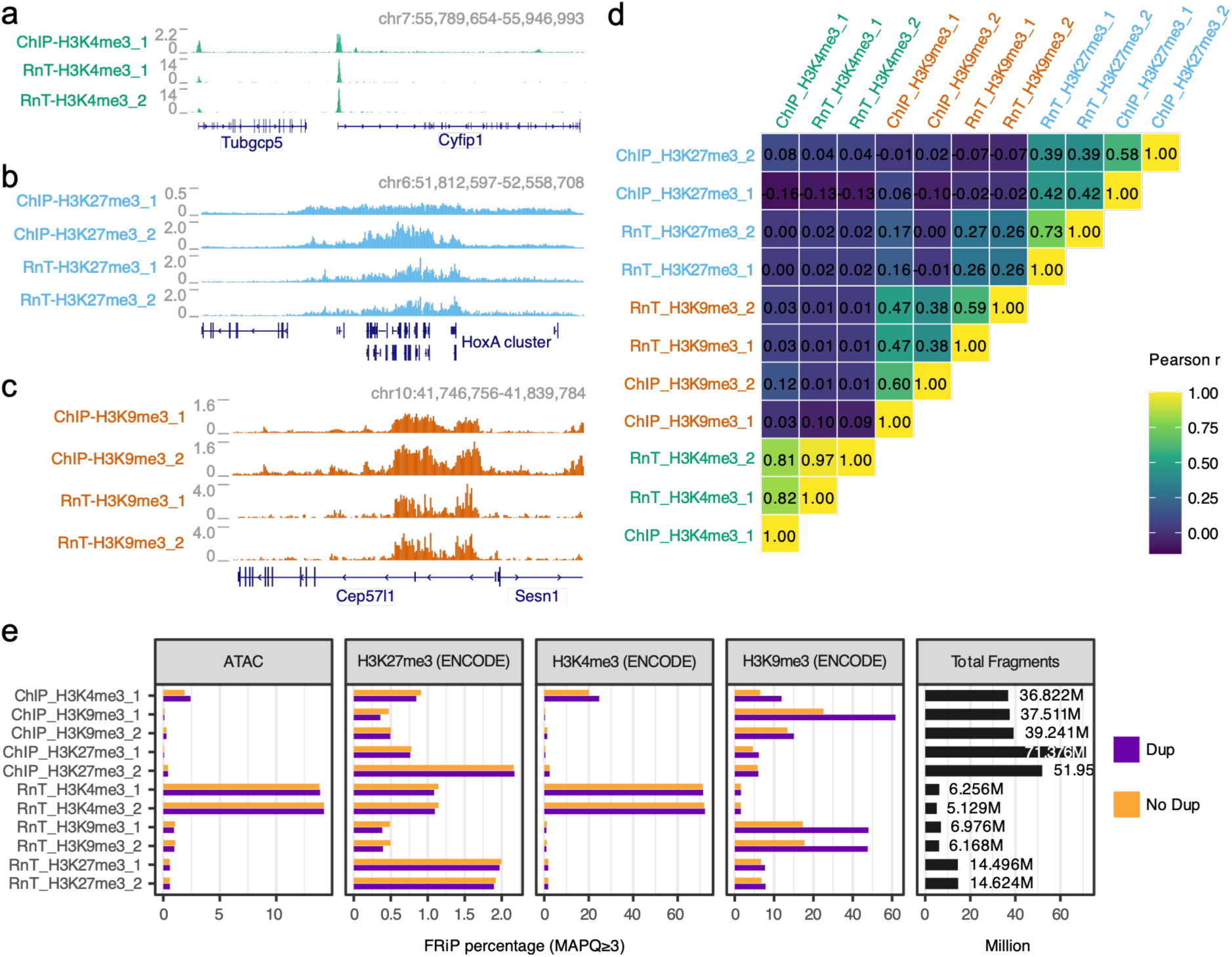
Read&Tag faithfully captures canonical histone modification patterns in frozen liver tissue. Read&Tag profiling of three histone modifications in archived mouse liver tissue and comparison to published ChIP-seq datasets using locus-level, genome-wide, and peak-based representations. **(a-c)** Genome browser tracks at representative loci showing profiles for H3K4me3 (A), H3K27me3 (B), and H3K9me3 (C) obtained from two frozen liver tissues using Read&Tag. Corresponding reference ChIP-seq tracks (refs. 43, 44) are shown for comparison, with gene annotations displayed below each locus. **(d)** Pairwise Pearson correlation heatmap comparing genome-wide signal distributions across Read&Tag biological replicates (n=2) and reference ChIP-seq datasets (refs. 43, 44) for the three histone modifications. Pearson correlations are shown for individual eCR-Tn5 configurations and replicates. **(e)** Fraction of reads in peaks (FRiP) for Read&Tag datasets targeting H3K27me3, H3K4me3, and H3K9me3. Bar plots show the percentage of sequencing fragments overlapping ENCODE reference peak sets, with duplicated (Dup) and non-duplicated (No Dup) reads indicated separately, and include comparisons to publicly available ChIP-seq datasets (refs. 43, 44).

Enrichment analysis demonstrated that Read&Tag captured substantial fractions of reads within ENCODE reference peak sets across all tested marks, with expected differences between focal and broad histone modifications and minimal sensitivity to PCR duplication or accessibility-driven transposition (Fig. 3e). To assess peak recovery in this tissue context, we performed precision-recall analysis using ENCODE ChIP-seq peak sets as references. Peaks identified using SEACR and GoPeaks showed high precision and intermediate recall for H3K4me3, whereas both metrics were reduced for the broader H3K27me3 and H3K9me3 domains (Supplementary Fig. 4a-b). These differences are consistent with the diffuse nature of broad histone modifications and variability in reference peak definitions across heterogeneous tissue samples. Together, these results demonstrate that Read&Tag enables specific and reproducible profiling of diverse histone modifications in long-term frozen tissue, capturing canonical chromatin features with high fidelity while requiring substantially less input material than conventional ChIP-seq approaches.

### Read&Tag enables histone modification profiling from low cell numbers

Given its robustness in frozen tissue, we next asked whether Read&Tag could be applied under conditions of limited cellular input. We generated Read&Tag profiles for H3K4me3, H3K27me3, and H3K9me3 from input samples of 500,000, 50,000, and 10,000 mESCs. Genome browser inspection showed that the characteristic genomic distributions of each histone modification were preserved across all input levels, with no apparent loss of signal at 10,000 cells (Fig. 4a). Consistent with these observations, pairwise correlation analysis revealed strong concordance between low-and high-input datasets for each histone mark, with profiles clustering primarily by modification rather than by input amount (Fig. 4b). Correlations were highest for the focal mark H3K4me3 and modestly reduced for the broader repressive marks at the lowest input, consistent with expected differences in domain structure and signal complexity. Aggregate signal profiles centered on GoPeaks-defined consensus peak regions generated from 500,000 cells further confirmed the preservation of mark-specific enrichment patterns at reduced input, including promoter-proximal enrichment for H3K4me3 and broad domain-level enrichment for H3K27me3 and H3K9me3 (Fig. 4c-e).

**Figure 4.**
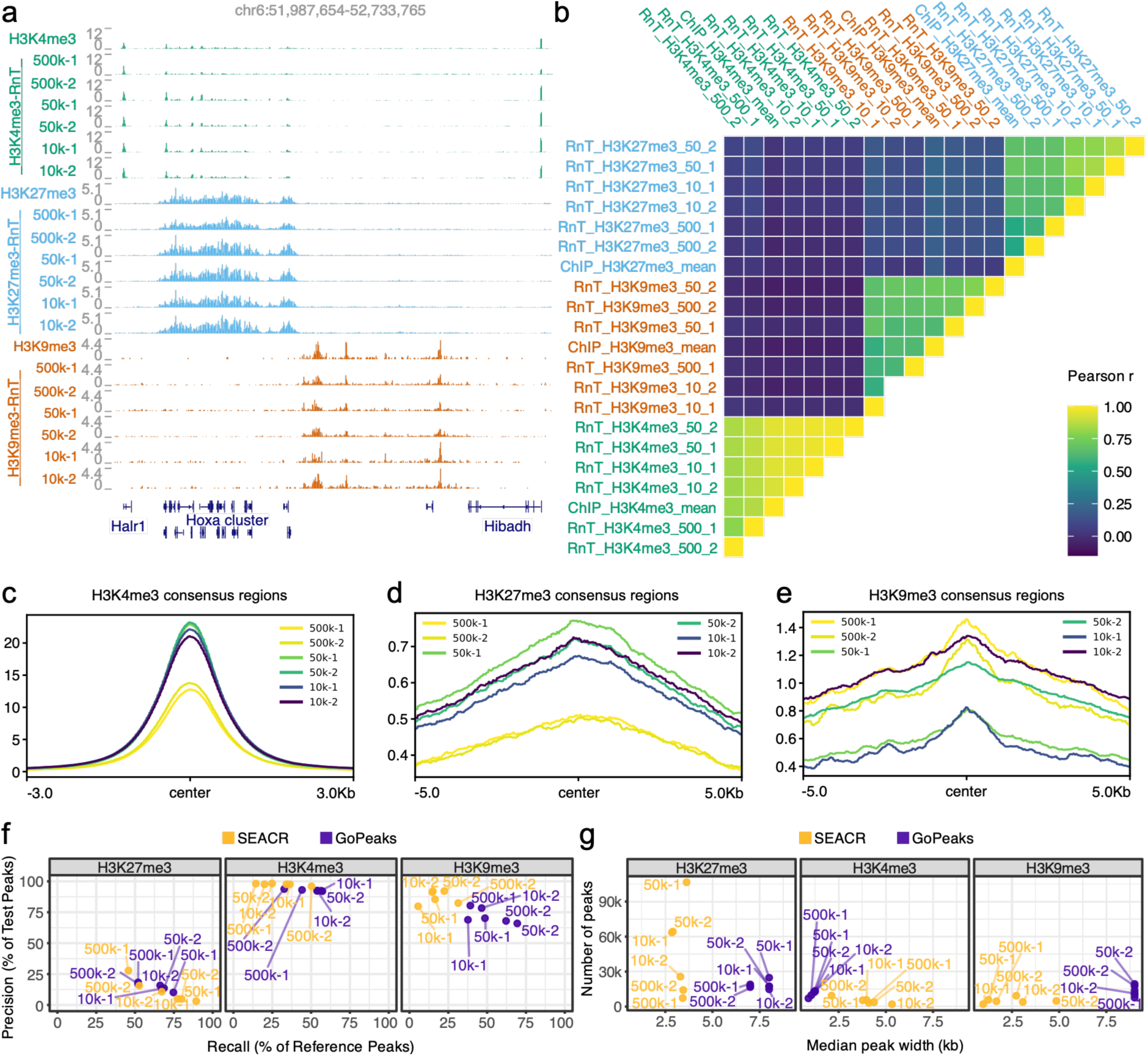
Histone modification profiling by Read&Tag at low cellular input. Performance of Read&Tag for profiling multiple histone modifications across decreasing input cell numbers, including representative genomic loci, genome-wide concordance, aggregate signal behavior, and peak-level benchmarking. **(a)** Genome browser tracks at a representative locus showing Read&Tag profiles for H3K4me3, H3K27me3, and H3K9me3 generated from 500,000, 50,000, and 10,000 input cells across biological replicates (n=2). Tracks are displayed alongside gene annotations to illustrate genomic signal distribution across input levels. **(b)** Pairwise Pearson correlation heatmap comparing genome-wide Read&Tag signal distributions across histone marks, reader configurations, input levels, and biological replicates, with color indicating correlation strength. **(c-e)** Aggregate Read&Tag signal profiles centered on GoPeaks-defined consensus peak regions for H3K4me3 (c), H3K27me3 (d), and H3K9me3 (e) generated from 500,000 cells. Curves represent signal intensity across peak-centered windows for each input level and biological replicate, illustrating signal behavior across decreasing cell numbers. **(f)** Precision-recall analysis comparing peak recovery across input levels for H3K27me3, H3K4me3, and H3K9me3. Each panel corresponds to a histone modification and displays individual datasets as points positioned by recall (x-axis; percentage of reference peaks recovered) and precision (y-axis; percentage of test peaks overlapping reference peaks). Points are colored by peak-calling method (SEACR or GoPeaks) and labeled according to input level and biological replicate. MACS2-defined reference peak sets are derived from ChIP-seq datasets and used as a common basis for comparison across conditions. **(g)** Summary of peak properties across input levels, showing the relationship between the number of identified peaks and median peak length for each histone mark and peak-calling strategy, enabling comparison of domain continuity and peak structure under reduced input conditions.

To quantify peak recovery across input levels, we performed precision-recall analysis using the ChIP-seq-defined (MACS2) consensus peaks as the reference set. GoPeaks achieved balanced precision and recall across histone marks and input levels, whereas SEACR showed higher precision but reduced recall for H3K4me3 and H3K9me3, consistent with its conservative peak-calling behavior (Fig. 4f). We next examined peak characteristics across input levels. Median peak width remained largely stable across all conditions, indicating consistent detection of both focal and broad domains under limiting input (Fig. 4g). GoPeaks identified comparable numbers of peaks across inputs and preserved broader peak widths for repressive histone marks, supporting robust domain-level signal recovery at low cell numbers.

Together, these results demonstrate that Read&Tag supports accurate histone modification profiling from low cell inputs, preserving genomic specificity and domain architecture across a broad dynamic range of input material.

### Read&Tag enables profiling of transcription-associated chromatin features via phospho-specific targeting of RNA polymerase II

Having established that Read&Tag enables robust chromatin profiling across diverse sample types and input regimes, we next asked whether the approach can be extended beyond histone modifications. For this, we adapted the method for profiling RNA polymerase II phosphorylated at serine 2 of the C-terminal domain (RNAP2-Ser2ph). We employed a phospho-specific RNAP2-Ser2ph probe ^45^ fused to Tn5, enabling direct targeting of elongating RNA polymerase II without reliance on conventional antibodies (Fig. 5a, Supplementary Fig. 5a).

**Figure 5.**
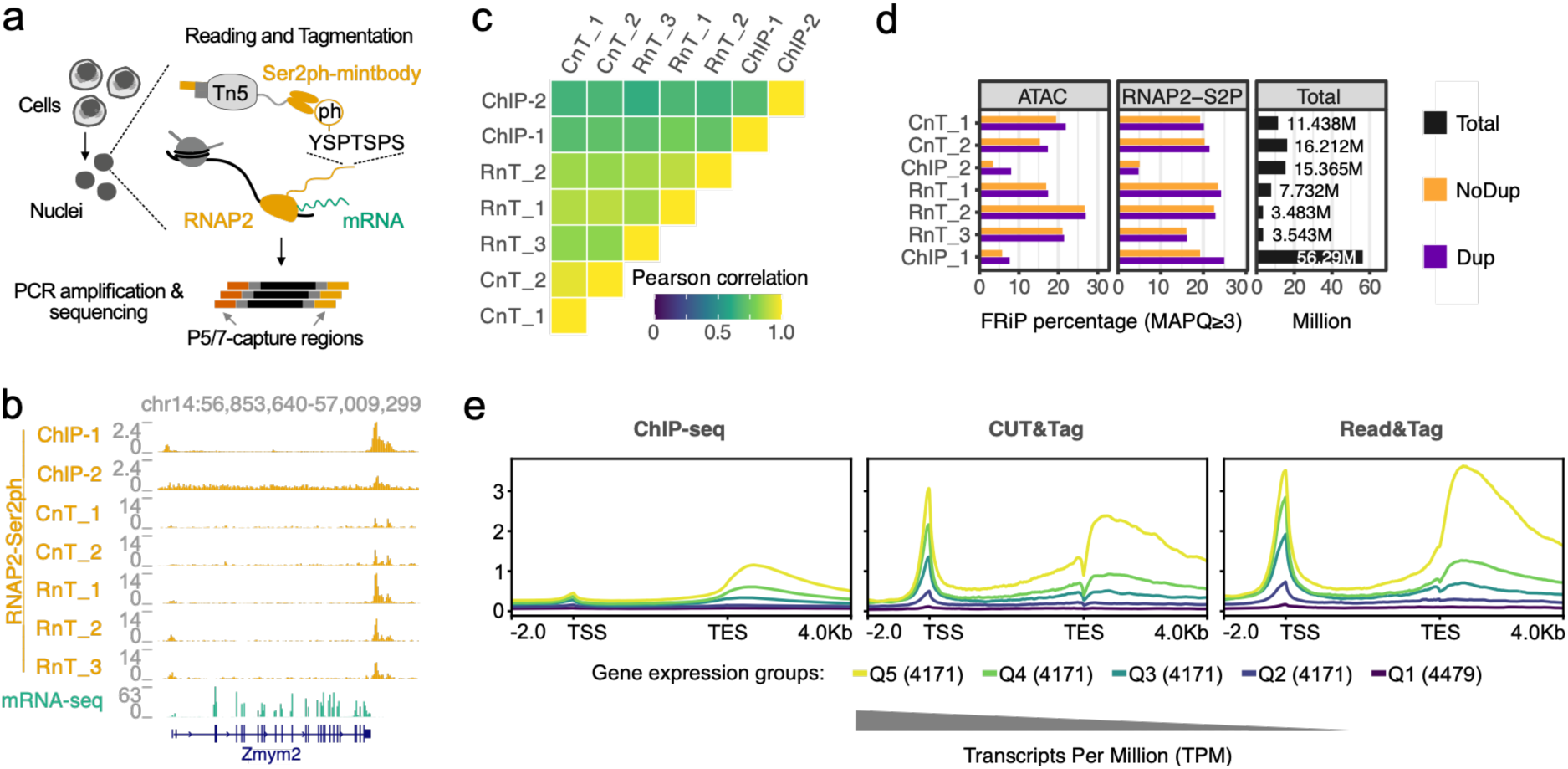
Read&Tag enables profiling of transcription-associated chromatin features via phospho-specific targeting of RNA polymerase II . Extension of Read&Tag to the profiling of a non-histone chromatin feature: RNA polymerase II phosphorylated at serine 2 of the C-terminal domain (RNAP2-Ser2ph). **(a)** Schematic of RNAP2-Ser2ph Read&Tag. A phospho-specific RNAP2-Ser2ph probe is coupled to Tn5 to enable targeting of elongating RNA polymerase II in permeabilized nuclei, generating barcoded sequencing libraries. **(b)** Genome browser tracks at representative loci showing RNAP2-Ser2ph signal obtained by ChIP-seq (refs. 46, 47), CUT&Tag (ref. 7), and Read&Tag, displayed together with mean mRNA-seq signal (n=4). CUT&Tag (n=2) and Read&Tag (n=3) signals are shown across biological replicates. **(c)** Pairwise Pearson correlation matrix comparing genome-wide RNAP2-Ser2ph signal distributions obtained by ChIP-seq, CUT&Tag, and Read&Tag across biological replicates. **(d)** Bar plots showing the fraction of sequencing reads overlapping ChIP-seq-defined peak sets (MACS2) for RNAP2-Ser2ph and accessibility-associated regions datasets generated by ATAC-seq. Duplicated and non-duplicated reads are indicated separately. **(e)** Metagene profiles of RNAP2-Ser2ph signal across the bodies of protein-coding genes (mm10), aligned at transcription start sites (TSS) and transcription end sites (TES). Genes are grouped by mRNA-seq expression level, with Q1 representing transcriptionally silent genes (TPM = 0) and Q2-Q5 representing equal-sized quantiles of expressed genes (TPM > 0). Profiles are shown for ChIP-seq (n=2), CUT&Tag (n=2), and Read&Tag mean signals (n=3) to illustrate transcription-associated signal patterns across expression strata.

Genome browser inspection showed that RNAP2-Ser2ph Read&Tag profiles closely recapitulated published RNAP2-Ser2ph ChIP-seq ^46,47^ and CUT&Tag ^7^ signal patterns at representative loci in mESCs, with enrichment over actively transcribed gene bodies and concordance with RNA-seq signal (Fig. 5b). Genome-wide correlation analysis further confirmed high agreement between RNAP2-Ser2ph Read&Tag and both ChIP-seq and CUT&Tag datasets, with reproducible correlations across biological replicates (Fig. 5c). FRiP analysis further showed that RNAP2-Ser2ph Read&Tag achieved substantial enrichment within ChIP-seq-defined peak regions, comparable to CUT&Tag, while exhibiting signal in accessible chromatin regions and characteristic nucleosomal fragment length distributions (Fig. 5d, Supplementary Fig. 5b). These features are consistent with targeted profiling of engaged RNA polymerase II rather than accessibility-driven transposition.

To assess whether Read&Tag captures transcriptional activity quantitatively, we performed metagene analysis by grouping protein-coding genes according to mRNA-seq expression levels. Q1 comprised transcriptionally silent genes (TPM = 0), and Q2-Q5 represented equal-sized quantiles of expressed genes (TPM > 0). RNAP2-Ser2ph Read&Tag signal was minimal over transcriptionally silent genes and increased progressively across expression quantiles Q2-Q5, with enrichment toward transcription end sites, consistent with accumulation of elongating RNA polymerase II and transcriptional activity (Fig. 5e). Notably, RNAP2-Ser2ph Read&Tag signal scaled with gene expression, indicating that Read&Tag captures graded and within-sample differences in transcriptional elongation rather than merely distinguishing active from inactive genes.

To determine whether phospho-specific RNAP2 Read&Tag performs comparably outside mouse ES cells, we applied the same RNAP2-Ser2ph reader-Tn5 fusion to human MCF7 cells. Genome browser tracks showed that Read&Tag reproduced the RNAP2-Ser2ph distribution obtained by two published ChIP-seq datasets ^48^ at representative loci, including transcribed gene bodies whose signal tracked mRNA-seq coverage (Supplementary Fig. 5c). Read&Tag replicates were reproducible genome-wide (Pearson r = 0.77-0.86), comparable to the agreement between the two ChIP-seq datasets (r = 0.82), and correlated with ChIP-seq at r = 0.56-0.69 (Supplementary Fig. 5d). At matched sequencing depth (13-16 million fragments, MAPQ≥3), Read&Tag recovered a two-to three-fold higher fraction of reads within ChIP-seq-defined RNAP2-Ser2ph peaks than ChIP-seq itself (∼16-20% versus ∼7-9%; Supplementary Fig. 5e). Metagene analysis in MCF7 cells recapitulated the behavior observed in mESCs: signal was near-background over silent genes and increased across expression quantiles, with pronounced accumulation toward transcription end sites, and this expression-dependent gradient was more clearly resolved by Read&Tag than by ChIP-seq (Supplementary Fig. 5f).

Together, these results demonstrate that Read&Tag can be extended beyond histone modifications to profile transcription-coupled chromatin features via phospho-specific targeting of elongating RNA polymerase II, providing a unified and modular framework for mapping both epigenetic and transcriptional features.

### Read&Tag with combinatorial engineered chromatin readers enables mapping of bivalent chromatin

Promoters of developmental genes carry both H3K4me3 and H3K27me3, a bivalent state thought to poise these genes for timely activation during differentiation ^49–51^. Mapping combinatorial chromatin states of this kind remains a major challenge, because the available approaches infer co-occurrence indirectly, either from separate single-mark experiments or from sequential immunoprecipitation. To address this, we engineered a combinatorial chromatin reader fused to Tn5 whose recruitment requires the simultaneous presence of H3K4me3 and H3K27me3 in the same chromatin fragment, enabling targeted tagmentation of bivalent chromatin (hereafter bivalent Read&Tag; Supplementary Fig. 6a).

Genome-wide profiles showed that the combinatorial reader localised preferentially to regions marked by both H3K4me3 and H3K27me3, whereas regions carrying either modification alone showed minimal enrichment. The monovalent H3K4me3-and H3K27me3-Read&Tag profiles were instead enriched at their respective single-mark regions and overlapped bivalent loci only partially (Fig. 6a). To systematically assess chromatin-state specificity, we projected 1-kb windows across the genome into a chromatin-state space defined by the percentile-ranked H3K4me3 and H3K27me3 ChIP-seq signals (Supplementary Fig. 6b). Monovalent Read&Tag signals were largely confined to their cognate single-mark axes, whereas the combinatorial reader was selectively enriched in the high-H3K4me3/high-H3K27me3 quadrant corresponding to bivalent chromatin (Fig. 6b). Among the top 1% of windows by signal, the combinatorial reader was strongly and selectively enriched in the bivalent state and correspondingly depleted from low-signal and monovalent states (Fig. 6c). Odds ratios computed over the same windows gave the same result (Supplementary Fig. 6c). Monovalent readers showed the reciprocal bias toward their related state, with H3K27me3-Read&Tag additionally enriched at bivalent regions, consistent with the scarcity of H3K27me3-only domains in mouse ES cells.

**Figure 6.**
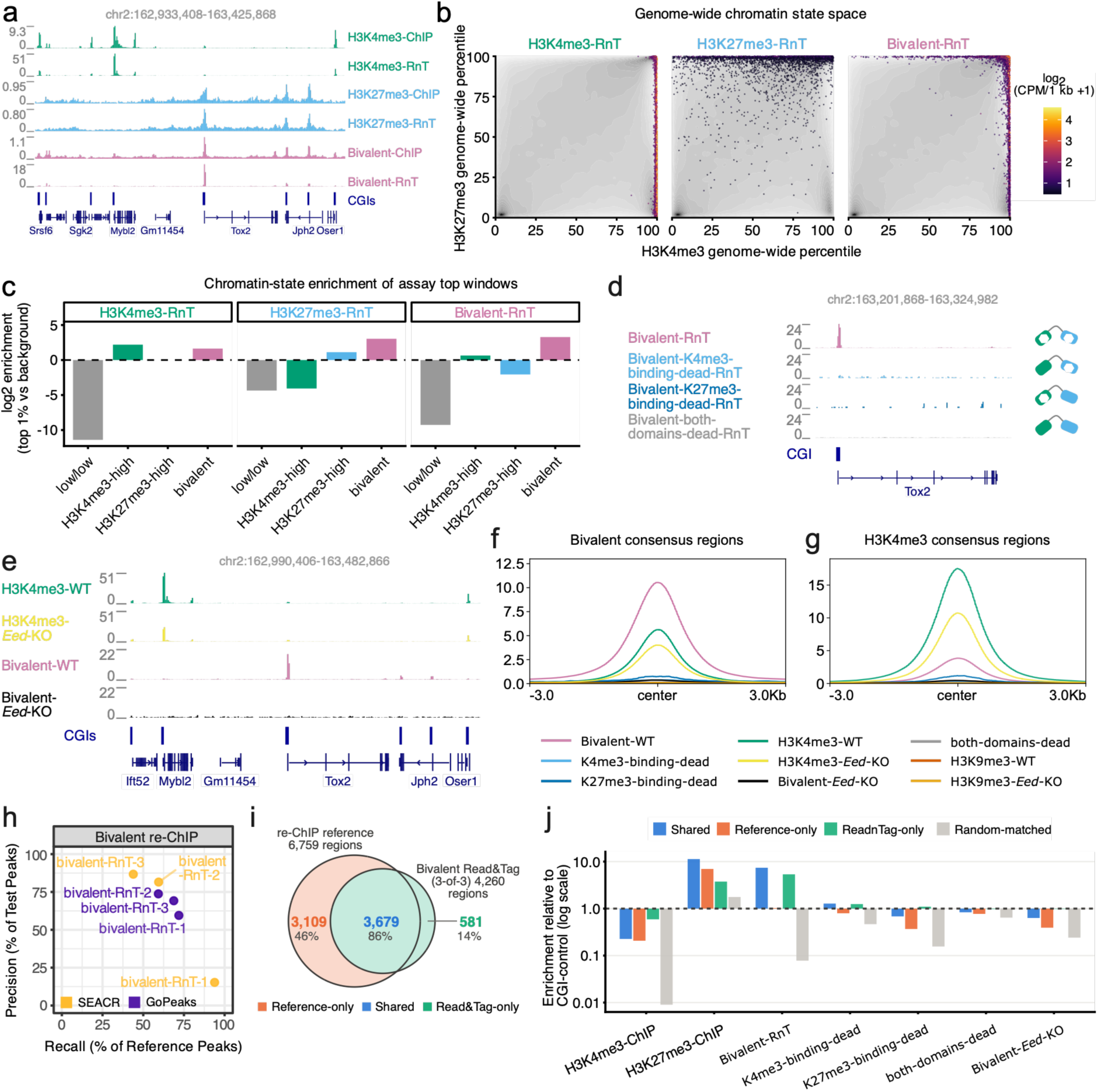
Read&Tag with combinatorial engineered chromatin readers enables mapping of bivalent chromatin. Generation and validation of an eCR-Tn5 fusion targeting bivalently H3K4me3-H3K27me3 modified chromatin. **(a)** Genome browser view at a representative locus showing combinatorial engineered chromatin reader (eCR) localization at H3K4me3-and/or H3K27me3-marked regions. Tracks: mean combinatorial Read&Tag (n=3), monovalent Read&Tag (n=2) and published bivalent-reader ChIP-seq (n=2, ref. 26) signals. Gene models and CpG islands (CGIs) below the tracks in (a), (d) and (e). **(b)** 1-kb genomic windows projected into chromatin-state space defined by percentile-ranked H3K4me3 (x) and H3K27me3 (y) ChIP-seq signal. Grayscale density, all windows; colored points, the top 1% windows per Read&Tag assay, shaded by signal (log2 CPM per 1 kb + 1). **(c)** Log2 enrichment of these top 1% windows over the genome-wide background in four chromatin states: low/low, H3K4me3-high, H3K27me3-high and bivalent. **(d)** As in (a), for combinatorial Read&Tag reader mutants in wildtype cells: combinatorial (Bivalent-RnT, n=3), H3K4me3-binding-domain dead (n=4), H3K27me3-binding-domain dead (n=4) and both-domains dead (n=3). **(e)** As in (a), in wildtype (WT) versus H3K27me3-deficient (Eed-KO) cells: H3K4me3 Read&Tag (WT, n=3; Eed-KO, n=2) and combinatorial Read&Tag (WT, n=3; Eed-KO, n=2). **(f-g)** Aggregate signal ±3 kb around consensus peak centers for combinatorial (Bivalent; f) and H3K4me3 (g) Read&Tag in wildtype cells, one curve per modality. **(h)** Precision-recall of peak recovery against a consensus reference of 6,759 bivalent regions from sequential ChIP (re-ChIP) in mouse ES cells (ref. 52). Each point, one dataset–peak-caller pair; precision, called peaks overlapping the reference; recall, reference peaks recovered (minimum overlap 100 bp). **(i)** Overlap between the 6,759 reference regions and the 4,260 bivalent regions called by combinatorial Read&Tag in all three replicates (≥1 shared bp; percentages relative to each set’s total). The shared area is labelled with the Read&Tag-side count (3,679, used in (j)); the reference-side count (3,650) is lower because some reference regions overlap more than one Read&Tag region. **(j)** Enrichment of each dataset over non-bivalent CpG islands: coverage (CPM per bp, pooled across all regions of a class) divided by the same at the 8,062 CGIs overlapping neither bivalent definition. Dashed line, signal indistinguishable from a non-bivalent CGI. Size-and chromosome-matched random regions serve as a null; H3K4me3-ChIP falls below one because bivalent regions carry less H3K4me3 than the average non-bivalent CGI.

To test whether targeting requires both reader-histone interactions, we introduced amino-acid substitutions that disrupt ligand binding by each reader module (TAF3 D890A/W891A; CBX2 W33A/W36A), individually or in combination, generating two single-dead-domain constructs and one both-domains-dead construct. To test in parallel whether targeting requires both modifications to be present on chromatin, we profiled *Eed*-knockout mouse ES cells, which lack H3K27me3 globally (Supplementary Fig. 6d). Genome browser tracks at representative loci showed that mutating either reader domain, mutating both, or removing H3K27me3 each strongly reduced combinatorial Read&Tag enrichment at regions carrying both modifications (Fig. 6d,e). Quantification across bivalent consensus regions supported this observation. Taking the both-domains-dead construct as the reference for untargeted tagmentation (and therefore as 1 by definition) enrichment was ∼39-fold for the intact reader but only ∼1.9-fold for the single-domain mutants and 2.1-fold in *Eed*-KO cells (Fig. 6f and Supplementary Table S4). Every construct yielded a library, so tagmentation itself was preserved, but the fragment-length distributions of the mutant constructs shifted from the nucleosomal pattern of the intact reader (Supplementary Fig. 6e) toward sub-nucleosomal fragments (Supplementary Fig. 6f,g), as expected for tagmentation that is no longer reader-guided.

The loss of signal was specific to the combinatorial reader. In *Eed*-KO cells, H3K4me3-Read&Tag remained enriched at H3K4me3 consensus regions, confirming that its target modification is retained in these cells, and H3K9me3-Read&Tag remained enriched at constitutive heterochromatin (Fig. 6g and Supplementary Fig. 6h,i). FRiP in *Eed*-KO cells was 14.7-29.6% for H3K4me3-Read&Tag at H3K4me3 regions and 21.6% for H3K9me3-Read&Tag at H3K9me3 regions, compared with 49% and 21.8% in wild-type cells (Supplementary Table S4). Thus, in cells that retain H3K4me3 but lack H3K27me3, the combinatorial reader loses essentially all enrichment while monovalent readers remain enriched at their target marks. Together with the reader-domain mutants, these results indicate that recruitment of the combinatorial reader, and hence reader-guided tagmentation, requires both intact histone-reader domains and the co-occurrence of H3K4me3 and H3K27me3 in the same chromatin region.

To benchmark recovery of bivalent chromatin, we compared combinatorial Read&Tag peaks with a consensus set of 6,759 bivalent regions defined by sequential ChIP-seq (re-ChIP) in mouse ES cells ^52^. Precision and recall depended strongly on the peak caller: GoPeaks was consistent across replicates (59-74% precision at 59-72% recall), whereas SEACR varied widely (15-87% precision at 44-94% recall; Fig. 6h). We therefore used GoPeaks calls supported by all three replicates, which raised precision to 86% at the cost of recall (54%) and defined 4,260 bivalent regions. Of these, 3,679 overlapped the reference set and 581 had no reference counterpart. On the reference side, 3,650 of the 6,759 regions were recovered and 3,109 were not (Fig. 6i). The two intersection counts differ because some reference regions overlap more than one Read&Tag region.

To characterize the three classes independently of Read&Tag, we used conventional H3K4me3 and H3K27me3 ChIP-seq and calculated the coverage in each class (CPM per bp, pooled across regions) relative to coverage at the 8,062 CpG islands that overlap neither bivalent set. A value of 1 therefore denotes signal indistinguishable from a non-bivalent CpG island. Because bivalent regions are predominantly CpG-island promoters^49,50^, this normalization removes what CpG islands have in common irrespective of their modification state. H3K4me3 values consequently fall below 1, indicating that bivalent regions carry less H3K4me3 than the average non-bivalent CpG island (Fig. 6j). The 3,109 missed reference regions carried less H3K27me3 than shared regions (7.0 versus 11.3) and no more H3K4me3 (0.21 versus 0.23). Of these, 1,666 were nevertheless called in at least one replicate and were lost to the three-replicate requirement, leaving 1,443 regions (21% of the reference set) undetected in every replicate (Supplementary Fig. 6j). The 581 Read&Tag-only regions carried less H3K27me3 than shared regions (3.8 versus 11.3) but were still nearly fourfold enriched over non-bivalent CpG islands, and carried 2.6-fold more H3K4me3 (0.59 versus 0.23), so neither modification alone predicted which regions were detected. At these regions the single-domain mutants, the both-domains-dead construct and the *Eed*-KO samples were all indistinguishable from control (0.98-1.25), as expected for a reader that requires both modifications. Together, these properties are consistent with bivalent regions that sequential ChIP-seq did not detect.

All of the above experiments were performed in mouse ES cells. To ask whether combinatorial targeting depends on that chromatin background, we applied the same combinatorial reader-Tn5 fusion to human MCF7 cells. Genome-wide chromatin-state analysis showed that the highest-signal windows again fell in the high-H3K4me3/high-H3K27me3 quadrant, the state-space signature of bivalent chromatin (Supplementary Fig. 6k). Collectively, these results demonstrate that combinatorial Read&Tag enables selective mapping of genomic regions marked by co-occurring H3K4me3 and H3K27me3 in mouse and human cells, and support engineered combinatorial readers as a general strategy for profiling co-occurring histone modifications *in situ*.

### Multifactorial profiling of epigenetic landscapes with Read&Tag

To determine whether Read&Tag can be extended to simultaneous profiling of multiple chromatin features within the same sample, we developed a multifactorial implementation (Multi-Read&Tag) targeting four marks: active promoter-associated H3K4me3; H3K27me3, associated with developmentally regulated Polycomb-repressed domains; H3K9me3, marking constitutive heterochromatin; and RNAP2-Ser2ph, indicative of transcriptional elongation (Fig. 7a). These targets span repressive and transcriptionally active chromatin environments, differ in genomic abundance, and range from focal promoter peaks to multi-kilobase domains.

**Figure 7.**
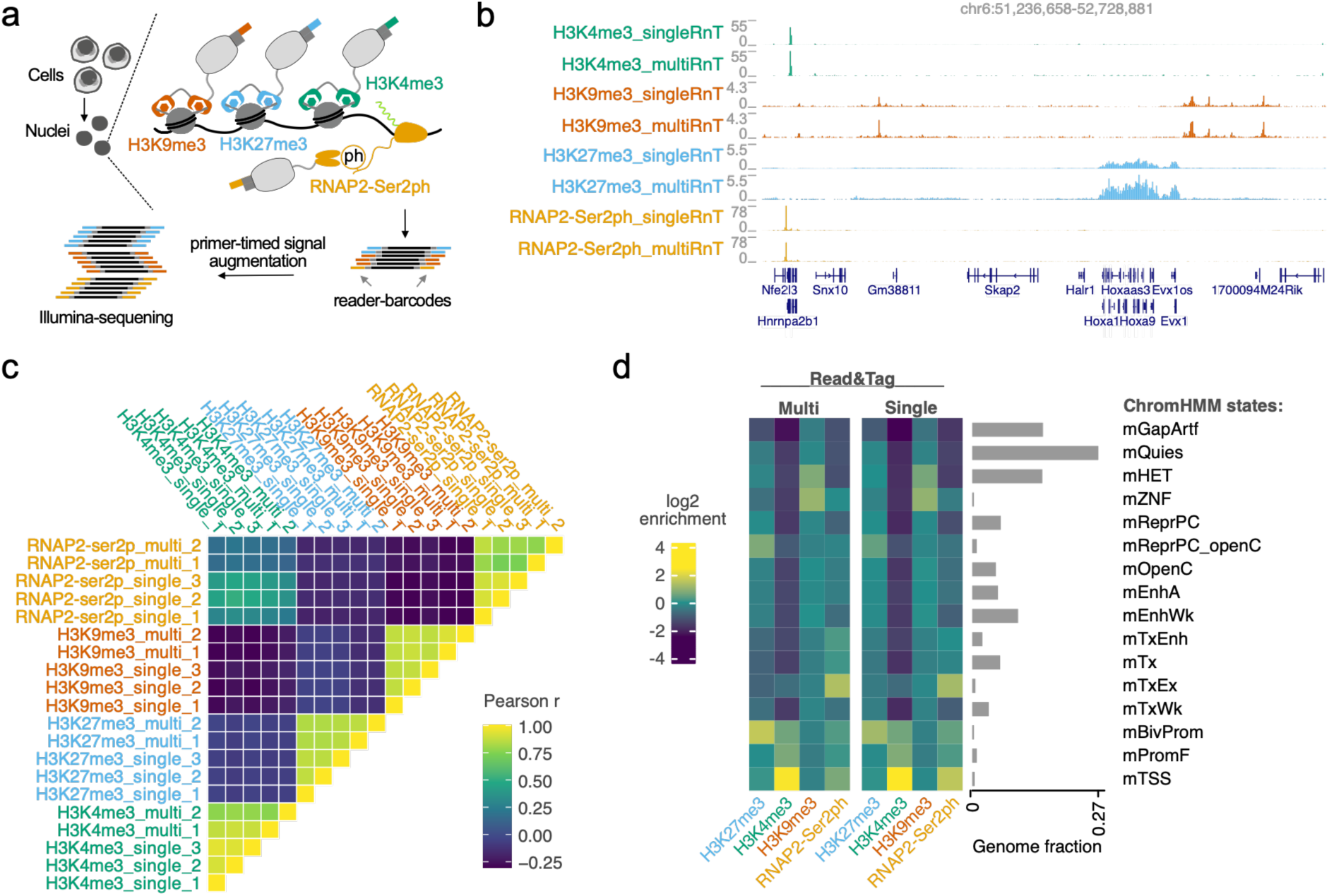
Simultaneous multifactorial chromatin profiling with Multi-Read&Tag. Simultaneous profiling of multiple, distinct chromatin features within the same Read&Tag experiment using barcoded reader-Tn5 fusions and primer-timed signal augmentation. **(a)** Schematic overview of the Multi-Read&Tag workflow. Reader-Tn5 fusion proteins targeting H3K4me3, H3K9me3, H3K27me3, and RNAP2-Ser2 phosphorylation are pre-loaded with reader-specific barcodes followed by simultaneous chromatin binding and tagmentation in permeabilized nuclei. Primer-timed signal augmentation (PTSA) is used during PCR amplification to balance the representation of targets with different genomic abundances by initiating amplification of individual targets at defined cycles. **(b)** Genome browser tracks showing representative loci for H3K4me3, H3K9me3, H3K27me3, and RNAP2-Ser2ph CPM-normalized signals profiled individually (“single”; average signal of three biological replicates) or simultaneously (“multi”; average signal of two biological replicates) using Multi-Read&Tag, illustrating target-specific signal patterns across conditions and replicates. **(c)** Pairwise Pearson correlation matrix restricted to the union of the four ChIP-seq peak sets (164,827 windows) comparing signal distributions at non-overlapping 1-kb windows for single-target Read&Tag and multi-target Read&Tag biological replicates across the four profiled features. Mean Pearson r, multifactorial versus single-target and single-target versus single-target respectively: H3K9me3 0.87/0.87; H3K27me3 0.84/0.86; H3K4me3 0.83/0.93; RNAP2-Ser2ph 0.79/0.91. **(d)** Heatmaps showing enrichment of average Read&Tag signal for H3K4me3, H3K9me3, H3K27me3, and RNAP2-Ser2ph across ChromHMM-defined chromatin states for single-target Read&Tag and Multi-Read&Tag. Bar plots indicate the genomic fraction of each mouse ChromHMM state.

Multi-Read&Tag was implemented using reader-specific barcoded Tn5 complexes applied simultaneously to permeabilized nuclei, so that all four targets experience the same nuclei, the same binding reaction and the same tagmentation. Because the four targets differ substantially in genomic abundance, uniform amplification would yield a library dominated by the most abundant mark. We therefore introduced primer-timed signal augmentation (PTSA), in which amplification of each barcoded sub-library is initiated at a defined PCR cycle, so that sub-libraries derived from low-abundance features such as RNAP2-Ser2ph receive additional amplification cycles relative to high-abundance histone marks while all targets remain within a single reaction (Fig. 7a). The PTSA schedule used for each sub-library is reported in Supplementary Table S5.

Genome browser inspection and signal quantification at quantiles of expressed genes or consensus peak sets revealed that Multi-Read&Tag profiles closely recapitulated the expected genomic distributions of each target and were highly concordant with corresponding single-target Read&Tag profiles (Fig. 7b). To assess relative distribution of signal across the genome, we computed 1-kb correlations restricted to ChIP-seq consensus peak sets. Restricted to the union of the four sets (164,827 windows), the twenty libraries resolved into four target-specific blocks, with negative correlation between mutually exclusive chromatin types and every multiplexed library falling within its own block. Within each target’s own consensus peaks, multifactorial libraries agreed with single-target libraries nearly as well as single-target replicates agreed with one another (Fig. 7c).

To test directly whether the within-target quantitation of Read&Tag survives multiplexing and PTSA, we repeated the analysis of Fig. 5e on the multiplexed data. RNAP2-Ser2ph signal measured by Multi-Read&Tag increased gradually across gene-expression quantiles, with the same rank order and metagene shape as in single-target Read&Tag and a comparable dynamic range between the highest and lowest expression quantiles (Supplementary Fig. 7a-b). H3K4me3 showed the same behavior at transcription start sites, while H3K27me3 and H3K9me3 showed the same inverse and flat relationships in both formats. Within-target quantitation is therefore retained, including in the RNAP2-Ser2ph sub-library, which receives one of the two largest numbers of amplification cycles (15 cycles; Supplementary Table S5).

Aggregate signal at each target’s consensus peak set further confirmed that multifactorial targeting Multi-Read&Tag does not compromise specificity in comparison to single-target Read&Tag (Supplementary Fig. 7c-d). Because H3K9me3 shares essentially no genomic territory with the other three targets, any signal crossing between its channel and the others must be leakage. We fitted each channel against independent CUT&Tag profiles of all four targets. *In silico* spike-ins showed that leaks of 0.5-3.5% out of the H3K9me3 channel, and 3.5-10% into it, would be detected in more than 95% of trials. Of twelve multiplexed measurements (six comparisons × two replicates), nine returned zero. The other three were below the detection limit, and the largest (0.80%) was not found in the matched replicate. Single-target Read&Tag and CUT&Tag libraries, which cannot leak between barcodes, gave values that multiplexing exceeded by at most 0.4 percentage points (Supplementary Table S6).

To assess whether multifactorial Read&Tag preserves higher-order chromatin context, we examined signal enrichment of each target modification across ChromHMM-defined chromatin states ^53^. Multi-Read&Tag profiles exhibited state-specific enrichment patterns that closely matched those observed in single-target experiments, with H3K4me3 and RNAP2-Ser2ph enriched at transcriptionally active states, H3K27me3 at Polycomb-repressed domains and bivalent promoters, and H3K9me3 at constitutive heterochromatin and zinc finger gene clusters (Fig. 7d).

Together, these results establish Multi-Read&Tag as a practical approach for simultaneous profiling of multiple epigenetic and transcription-associated features within a single experiment, including features that genuinely co-occur in the genome. By combining modular barcoded eCR-Tn5 targeting with primer-timed signal augmentation, Multi-Read&Tag reproduces the genomic distribution, peak-level enrichment, chromatin-state specificity and within-target quantitative response of single-target Read&Tag, with no detectable exchange of signal between channels.

## Discussion

In this study, we introduce Read&Tag, a modular chromatin-profiling platform that leverages engineered chromatin readers fused to Tn5 transposase to enable specific, combinatorial, and multifactorial epigenomic profiling. By replacing antibodies with defined chromatin-binding domains, Read&Tag provides a flexible and extensible alternative to antibody-dependent in situ chromatin-profiling approaches, while retaining the sensitivity and workflow of transposase-based library preparation.

A central advantage of Read&Tag is its reader-guided specificity, which distinguishes it from both classical ChIP-seq and antibody-tethered tagmentation methods. Antibody-based methods such as ChIP-seq ^15^, ChIL-seq ^3^, CUT&RUN ^2^, CUT&Tag ^4^, and transposase-related approaches ^5–12^ rely on antibody affinity and specificity, requiring extensive optimization. In contrast, Read&Tag decouples chromatin targeting from antibody performance by employing recombinant reader domains with defined binding specificities, enabling more controlled and reproducible targeting of chromatin features. Because recognition is performed by a defined chromatin reader domain rather than by an affinity reagent, the targeting module itself becomes a design variable: its valency can be varied, its binding pocket mutated to provide a matched negative control, and two specificities combined in one complex so that recruitment requires the co-occurrence of both modifications. This combinatorial recognition ability is not available to antibody-or nanobody-tethered approaches, in which co-occurrence can only be inferred by intersecting independently measured modifications.

Read&Tag also addresses a known limitation of Tn5-based in situ methods: accessibility-driven background ^18^. While Tn5-based methods such as ATAC-seq exploit open chromatin bias, this property can confound profiling of chromatin states. Our analyses show that Read&Tag signal localization is predominantly driven by reader-chromatin interactions rather than accessibility, as evidenced by loss of signal upon reader mutation, low enrichment at accessible regions for repressive marks, and preservation of specificity across frozen tissue and low-input conditions. This feature is particularly advantageous for profiling broad repressive domains such as H3K27me3 and H3K9me3.

Beyond histone modifications and DNA methylation, Read&Tag can be readily extended to non-histone chromatin-associated features. By targeting elongating RNA polymerase II via a phospho-specific RNAP2-Ser2ph binder ^45^, we show that Read&Tag captures graded transcriptional activity across gene expression levels. The quantitative relationship between RNAP2-Ser2ph signal and mRNA abundance indicates that Read&Tag can effectively profile transcription-coupled chromatin features, highlighting its potential for integrated epigenomic-transcriptional analyses.

A key conceptual advance enabled by Read&Tag is the direct interrogation of combinatorial chromatin states. Using engineered combinatorial readers, we demonstrate selective enrichment of genomic regions co-marked by H3K4me3 and H3K27me3, capturing a chromatin feature that is not readily accessible with conventional antibody profiling. This capability underscores the potential of engineered chromatin readers to move beyond single-mark detection toward direct measurement of combinatorial chromatin states. While reader mutagenesis and loss of H3K27me3 in EED-deficient cells establish that recruitment of the combinatorial reader strictly requires both modifications and both binding surfaces, our data do not formally exclude engagement of two adjacently positioned bivalently modified nucleosomes. Definitive assignment will require *in vitro* reconstitution on defined, asymmetrically modified mono- and di-nucleosomes.

Building on Read&Tag’s modularity, we further establish Multi-Read&Tag as a strategy for simultaneous profiling of multiple chromatin features within a single experiment. Using differentially barcoded eCR-Tn5 complexes and controlled signal balancing via primer-timed signal augmentation, we demonstrate simultaneous profiling of four distinct chromatin features (H3K4me3, H3K27me3, H3K9me3, and RNAP2-Ser2ph) without detectable cross-contamination or loss of specificity. Compared to multi-CUT&Tag ^7^, ACT2-seq ^8^, MulTI-Tag ^10^ and nanobody-based multimodal approaches ^11,12^, Multi-Read&Tag offers a complementary strategy that does not require primary antibodies, secondary nanobodies, or genetic fusion of epitopes to the target ^13,14^, and is therefore applicable to endogenous chromatin modifications in unmodified cells and in primary material.

Despite these advantages, several limitations should be considered. As with other Tn5-based approaches, Read&Tag may be influenced by chromatin accessibility under certain conditions, and highly compact heterochromatin may remain challenging to profile at very low input. We also recommend including an exogenous chromatin spike-in for perturbations expected to alter global modification levels more severely. Because primer-timed signal augmentation equalizes target representation by applying different numbers of amplification cycles per target, Multi-Read&Tag does not preserve relative target abundance. In addition, the current repertoire of high-affinity, modification-specific reader domains and probes is more limited than the available antibody collections, potentially restricting the range of immediately accessible targets. However, ongoing advances in protein engineering ^54^, directed evolution ^27,55^, and protein *de-novo* design ^56–60^ are likely to rapidly expand the available repertoire and facilitate further optimization of selectivity and affinity.

In summary, Read&Tag provides a modular, antibody-independent method for chromatin profiling that unifies epigenetic and transcription-associated modifications within a single experimental setup. By combining recombinant eCR-Tn5 fusions with flexible barcoding and amplification strategies, Read&Tag enables robust profiling across diverse sample types, input regimes, and chromatin features. We anticipate that this approach will be broadly applicable to studies of development and disease, and will serve as a foundation for future extensions toward single-cell implementations, additional chromatin-associated modalities, testing of chromatin-targeting drugs, and simultaneous multi-omics measurements.

## Methods

### Generation of reader-Tn5 constructs

pTXB1-Tn5 was a gift from Rickard Sandberg (Addgene plasmid # 60240; http://n2t.net/addgene:60240; RRID:Addgene_60240) ^61^. PB533-42B3mutAC2-sfGFP was a gift from Hiroshi Kimura (Addgene plasmid # 186777; http://n2t.net/addgene:186777; RRID: Addgene_186777) ^45^. Engineered chromatin reader and mintbody sequences were synthesized at Twist Bioscience with 30 nt overhangs (5’-gaattgtgagcggataacaattcccctctagaaataattttgttaactttaagaaggagatatacat-3’; 5’-catatgattaccagtgcactgcatcgtgcg-3’) and cloned into XbaI and NdeI linearized pTXB1-Tn5 via Gibson assembly. The full sequences of the fusion proteins and oligonucleotides are available in the Supplementary Tables S2 and S3, respectively. All constructs used in this study have been deposited with Addgene (hold for publication) and available on request under an MTA in the meantime.

### Purification of reader-Tn5 fusions

pTXB1-reader-Tn5 fusion proteins were similarly purified as described in Picelli et al. ^61^ and Kayar-Okur et al. ^4^. The pTXB1-reader-Tn5 plasmid was transformed into T7 Express *lys^Y^/I^q^* competent *E. coli* cells (NEB, Cat.No. C3013I) following the manufacturer’s protocol. Single colonies were picked and analyzed using colony PCR with primers T7fw and Tn5_rv (Supplementary Table S3). Positive clones were inoculated into 30 mL LB medium (Serva, Cat.No. 48501.01) containing ampicillin (Carl Roth, Cat.No. K029.2) and grown overnight at 37 °C, shaking at 180 rpm. The next day, 2 L LB medium supplemented with 100 µg/mL carbenicillin (Carl Roth, Cat.No. 6344.4) and 2 % glucose (Millipore, Cat.No. 1.08342.1000) was inoculated with the entire overnight culture. Bacteria were grown to OD600 0.6 and chilled on ice for 60 min. Then, protein expression was induced with 0.25 mM Isopropyl β-d-1-thiogalactopyranoside (IPTG, Genaxxon bioscience, Cat.No. M3198.0025) at 16-18 °C on a shaker overnight. Bacteria expressing the Zinc-finger proteins, such as TAF3-PHD-domain containing constructs, were additionally treated with 0.2 μM ZnCl₂ during induction. Bacteria were harvested by centrifugation at 4000 x g at 4 °C for 10 min and resuspended in 80 mL of Wash buffer (20 mM HEPES-KOH pH 7.2, 0.8 M NaCl, 10 % glycerol, 0.2 % Triton X-100) containing 1 mM phenylmethylsulfonyl fluoride (PMSF, Carl Roth, Cat.No. 6367.2) and protease inhibitor cocktail (PIC) at 1x concentration (100x PIC made from EDTA-free protease inhibitor tablets; Roche, Cat.No. 05056489001). The bacterial suspension was lysed by sonication (10 cycles, 30 sec ON/60 sec OFF, 40 % duty cycle). The lysate was then cleared by centrifugation at 17371.6 x g for 10 min at 4 °C. Next, 2.1 mL of 10 % neutralized polyethylenimine (PEI, Merck, Cat.No. P3143-100ML) was added dropwise to the lysate to precipitate nucleic acid contaminants. This was followed by another centrifugation at 17371.6 x g for 10 min at 4 °C. A disposable column (Bio-Rad, Cat.No. 7321010) was packed with 2.5 mL of chitin slurry resin (New England Biolabs, Cat.No. S6651S), washed twice with 10 mL ddH_2_O, once with 10 mL Wash buffer, and then resuspended in the nucleic acid-depleted lysate. Proteins were bound for 40 min at 4 °C during continuous rotation on the chitin resin. Subsequently, the chitin resin was transferred back to the column. After the unbound lysate was drained, the resin was washed four times with 25 mL of Wash buffer. Next, 15 mL Elution buffer (20 mM HEPES-KOH pH 7.2, 0.5 M NaCl, 10 % glycerol) containing 100 mM dithiothreitol (DTT, Carl Roth, Cat.No. 6908.2) was added to the column. After letting 8 mL of Elution buffer drain from the column, the chitin resin was resuspended in the remainders of Elution buffer and incubated for 48 h at 4 °C with continuous rotation to facilitate intein-mediated cleavage from the resin. For Zinc-finger proteins, the elution buffer was additionally supplemented with 0.1 μM ZnCl₂. The eluate was collected and concentrated using an Amicon Ultra-4 Centrifugal Filter Unit 10 K (Merck, Cat.No. UFC9010) to a final volume of 0.3-0.5 mL. Subsequent dialysis was performed against a Dialysis buffer (20 mM HEPES-KOH pH 7.2, 0.5 M NaCl, 20 % glycerol). Protein concentration was determined by measuring UV-Vis absorbance at 280 nm, with extinction coefficients calculated using the Benchling biology software (https://benchling.com, 2026). The dialyzed protein was preserved for long-time storage by adding both Dialysis buffer and 80 % sterile glycerol (Sigma Aldrich, Cat.No. 15523-2.5L-M) to achieve a final concentration of 5.5 µM protein in 50 % glycerol. Protein stocks were snap-frozen in liquid nitrogen as 100 µL aliquots and stored at -70 °C.

### Adaptor loading of reader-Tn5 fusions

All Oligonucleotides used for adaptor loading were purchased from Microsynth AG (Balgach, Switzerland) and dissolved in Annealing buffer (10 mM Tris-HCl pH 8.0, 50 mM NaCl, 1 mM EDTA) at 200 µM final concentration and stored at -20 °C. Reader-Tn5 fusion proteins were loaded either with generic or reader-specific barcoded Tn5MEDS-A and Tn5MEDS-B adaptors. For Tn5MEDS-A generic or barcoded adaptors, 5 µL of each forward oligonucleotide (200 μM) was mixed with 5 µL of 5’ phosphorylated Tn5_Rev oligonucleotide (200 µM) to achieve equimolar amounts of oligonucleotides. Similarly, generic or barcoded Tn5MEDS-B are produced by mixing 200 μM forward oligonucleotide (5 μL) with an equimolar amount of 200 μM of 5’ phosphorylated Tn5_Rev oligonucleotide (5 μL). For a complete list of adaptor oligonucleotides, see Supplementary Table S3. Oligonucleotides were denatured at 95 °C for 5 min and then gradually cooled to facilitate annealing. 8 µL of preannealed Tn5MEDS-A/Tn5_Rev and 8 µL Tn5MEDS-B/Tn5_Rev oligonucleotide were mixed with 100 µL of 5.5 µM reader-Tn5 fusion protein. The mixture was incubated on a rotating platform for 1 h at room temperature (RT) and then stored at -20 °C.

### Cell culture

mESCs (HA36CB1, 129xC57BL/6)^62^, PRC1-degron mESCs^38^ (kindly provided by Rob Klose) and *Eed*-knockout mESCs^26^, were cultured on 0.2 % gelatine-coated dishes in mESC medium containing DMEM (Gibco, Cat.No. 31966-021) supplemented with 15 % FBS (PAN Biotech, Cat.No. P30-3030M), 1x non-essential amino acids (Sigma-Aldrich, Cat.No. M7145), 50 μM β-mercaptoethanol (Gibco, Cat.No. 31350-010), and homemade leukemia inhibitory factor (LIF). Cells were incubated at 37 °C in a humidified environment with 5 % CO₂. The culture medium was changed every 24 h, and cells were passaged every 2 days to ensure optimal growth conditions. To induce AID-RING1B degradation, PRC1-degron cells were treated with 500 µM auxin (IAA, Sigma-Aldrich, Cat.No. I3750; dissolved in water) in mESC medium for 8 or 24 h before collection with TrypLE (Gibco, Cat.No. 12605-028). Untreated cells served as the 0 h control. Human MCF-7 cells (kindly provided by Andreas Ladurner) were cultured on 0.2 % gelatine-coated dishes in medium containing DMEM (Gibco, Cat.No. 31966-021) supplemented with 10 % FBS (PAN Biotech, Cat.No. P30-3030M) and 1 % penicillin-streptomycin (Sigma-Aldrich, Cat.No. P4333). Cells were incubated at 37 °C in a humidified environment with 5 % CO₂ and passaged every 2-3 days. *Drosophila melanogaster* S2 cells (kindly provided by Peter Becker), used as spike-in chromatin, were cultured at 26 °C in Schneider’s medium (Thermo Fisher Scientific, Cat.No. 21720024) supplemented with 10 % FBS (PAN Biotech, Cat.No. P30-3031, or Capricorn Scientific, Cat.No. FBS-12A) and 1 % penicillin-streptomycin (Sigma-Aldrich, Cat.No. P4333). All cell lines tested negative for mycoplasma.

### Histone extraction and immunoblotting

Histones were acid-extracted as described^38^. Cell pellets (∼80 % confluent 10 cm plate) were washed in PBS, snap-frozen and stored at −80 °C. Thawed pellets were washed in 1 ml RSB (10 mM Tris-HCl pH 7.4, 10 mM NaCl, 3 mM MgCl₂, 20 mM N-ethylmaleimide), pelleted (240 × *g*, 5 min) and lysed in RSB with 0.5% NP-40 for 10 min on ice. Nuclei were pelleted (500 × *g*, 5 min), resuspended in 200 µl 5 mM MgCl₂ and extracted with 0.4 M HCl (final) for 20 min on ice. After centrifugation (18,000 × *g*, 20 min), histones in the supernatant were precipitated with 25% (v/v) TCA for 30 min on ice and pelleted (18,000 × *g*, 15 min). The pellet was washed twice with ice-cold acetone, resuspended in ddH₂O and quantified with the Qubit Protein Assay (Invitrogen, Q33212). All centrifugations were done at 4 °C.

0.5-3 µg of histones were denatured in 1× SDS loading buffer (from 5×: 10% SDS, 250 mM Tris pH 6.8, 0.5 M DTT, 50% glycerol, 0.05% bromophenol blue) at 95 °C for 5 min. They were separated on 15% SDS-PAGE gels in Tris-glycine-SDS running buffer (25 mM Tris, 192 mM glycine, 0.1% SDS) and wet-transferred to PVDF (Millipore, 0.45um, Cat. No.IPFL00010) in 48 mM Tris, 39 mM glycine, 1.3 mM SDS, 20% methanol (30 V, overnight, on ice). Membranes were blocked in 3% BSA/TBS for 1 h at RT. They were then incubated overnight at 4 °C with primary antibodies in 3% BSA/TBS-T: anti-H2AK119ub1 (Cell Signaling, Cat. No. 82405, 1:5000), anti-H3K27me3 (Diagenode, Cat. No. C15410195, 1:2000) and anti-H3 Abcam, Cat. No. ab10799, 1:2000; loading control). This was followed by IRDye 680RD/800CW secondary antibodies (LI-COR, Cat. No. 680 -anti Ms: 926-68070 / 800 -anti Rb: 926-32211, both 1:10,000) in TBS-T (0.1% Tween 20) for 1 h at RT. Each antibody step was followed by three 5-min TBS-T washes. Signals were acquired on an Odyssey CLx (LI-COR).

### Preparation of cell nuclei from cell lines

Cells were harvested, counted, and centrifuged at 600 x g for 3 min at RT, then washed twice with ice-cold 1x PBS. Cells were resuspended in 10 mL of nuclear extraction buffer (20 mM HEPES-KOH pH 7.9, 10 mM KCl, 0.5 mM spermidine (Sigma Aldrich, Cat.No. 85558-1G), 0.1 % Triton X-100, 20 % glycerol, 1x PIC) and incubated on ice for 10 min. Nuclei were pelleted at 1300 x g for 4 min at 4 °C. For native nuclei, the pellet was resuspended in ice-cold Read&Tag wash buffer (20 mM HEPES pH 7.5, 150 mM NaCl, 0.5 mM spermidine, 1x PIC) and used for Read&Tag without further treatment. To obtain fixed nuclei, the pellet was resuspended in 10 mL ice-cold 1x PBS and subsequently crosslinked by adding 62 µL of 16 % formaldehyde (final concentration 0.1 %; Thermo Fisher Scientific, 10 mL ampules, Cat.No. 28908) for 5 min at RT. The fixation was quenched by adding 266 µL of 2.5 M glycine (final concentration 65 mM; Serva, Cat.No. 23390.04). Fixed nuclei were centrifuged again at 1300 x g for 4 min at RT. The pellet was then resuspended in ice-cold Read&Tag wash buffer. Multi-Read&Tag was performed using fixed nuclei. All other experiments were done with native material. Prepared nuclei were either used immediately or cryopreserved with addition of dimethyl sulfoxide (DMSO; final concentration 10 %; Sigma-Aldrich, Cat.No. D2438-50ML). Cryopreservation was achieved using a controlled-rate freezing approach with gradual cooling (∼1 °C/min) using a Mr Frosty Freezing Container (Thermo Fisher Scientific, Cat.No. 5100-0001) before long-term storage at -70 °C.

### Read&Tag

Concanavalin A (Con A)-coated magnetic beads (Polysciences, BioMag(R)Plus Concanavalin A, Cat.No. 86057-3) were activated by two washes. 15 µL ConA-coated beads were used for each sample of 500,000 nuclei; 10 µL for each sample of 10,000-50,000 nuclei. For the washes, Binding buffer (20 mM HEPES-KOH pH 7.9, 10 mM KCl, 1 mM CaCl₂, 1 mM MnCl₂) was used at 10x the starting volume of ConA-coated beads. After, the ConA-coated beads were resuspended in the original volume of Binding buffer.

Per sample, 10,000-500,000 nuclei were aliquoted and adjusted to 1 mL with Read&Tag wash buffer. Activated ConA-coated beads were added and the mix incubated for 15 min at RT with continuous rotation to facilitate nuclei binding. After removing the unbound supernatant, nuclei-bound beads were resuspended in 1 mL of Read&Tag lysis buffer (20 mM HEPES pH 7.5, 150 mM NaCl, 0.5 mM spermidine, 2 mM EDTA, 1x PIC, 0.2 % Triton X-100, 0.05 % digitonin (Millipore, via Scientific Laboratory Supplies, Cat.No. 300410-1GM), 0.1 mg/mL recombinant Albumin (New England Biolabs, Cat.No. B9200S)) and permeabilized overnight at 4 °C with continuous rotation. Beads were washed three times in 0.9 mL of Read&Tag high-salt wash buffer (20 mM HEPES pH 7.5, 300 mM NaCl, 0.5 mM spermidine, 0.01 % digitonin, 1x PIC). Up to eight samples were processed together until this step; during the final wash, the nuclei-bound beads were split into individual samples and subsequently resuspended in 250 µL of Read&Tag high-salt buffer (20 mM HEPES pH 7.5, 300 mM NaCl, 0.5 mM spermidine, 0.01 % digitonin, 1x PIC, 0.25 % Triton X-100). For single-target mapping, 1-2 fmol of generic adaptor-loaded reader-Tn5 was added to the respective sample; for multiplexed mapping, a mixture of several 1-2 fmol barcoded adaptor-loaded reader-Tn5‘s were used. Binding was carried out for 1 h and 45 min at 4 °C with continuous rotation. Beads were washed four times for 5 min in Read&Tag high-salt buffer to eliminate unbound reader-Tn5. Tagmentation was started by resuspending the beads in 300 µL of Tagmentation buffer (20 mM HEPES pH 7.5, 300 mM NaCl, 10 mM MgCl₂, 0.5 mM spermidine, 0.01 % digitonin, and 1x PIC) and incubation at 37 °C with shaking at 450 rpm for 1 h. Tagmentation was stopped by adding EDTA (final concentration 16 mM), SDS (final concentration 0.1 %), and Proteinase K (final concentration 0.16 mg/mL), followed by incubation at 55 °C with shaking at 550 rpm for 1 h. Beads were separated magnetically, and the supernatant was collected for DNA purification. DNA was purified by adding 400 µL AMPure buffer (1.25 M NaCl, 20 % PEG 8000) and 100 µL SPRI paramagnetic beads (AMPure XP beads by Beckman Coulter, Cat.No. A63880; alternatively, MAGFLO NGS beads by Integra, Cat.No. 7002). The reaction was incubated for 15 min at RT, washed twice with 80 % ethanol (Sigma Aldrich, Cat.No. 32221-2.5L-M), air-dried, and eluted in 25-40 µL of 0.1x TE (Tris-EDTA; 1 mM Tris-HCl pH 8.0, 0.1 mM EDTA) buffer to obtain tagmented DNA.

### Calibrated Read&Tag

Spike-in calibration was performed using PRC1-degron mESC nuclei combined with S2 nuclei at a ratio of 10:1, immediately prior to ConA binding, permeabilisation, and tagmentation within the same reaction. Read&Tag was then performed as described above, using the H3K27me3 reader at 0, 8 and 24 hours after auxin treatment (two biological replicates were measured for each time point).

### Library preparation for single-target Read&Tag

To amplify Read&Tag libraries, 21 µL of tagmented DNA was mixed with 2 µL of uniquely barcoded i5 and i7 index oligonucleotides (10 µM each; for a full list of indexing oligonucleotides, see Supplementary Table S3) and 25 µL of NEBNext HiFi 2x PCR Master Mix (New England Biolabs, Cat.No. M0541S). The samples were placed in a thermocycler with a heated lid and enriched using the following program: 72 °C for 5 min (gap filling); 98 °C for 30 sec; 14-16 cycles of 98 °C for 10 sec and 65 °C for 45 sec; final extension at 72 °C for 5 min, and eventual cooling to 4 °C. Libraries were purified using SPRI paramagnetic beads (AMPure XP beads by Beckman Coulter; alternatively, MAGFLO NGS beads by Integra). 65 µL of SPRI paramagnetic beads were pipetted into the PCR reaction, thoroughly mixed, and incubated for 7 min at RT. Beads were washed twice with 80 % ethanol, air-dried for 2 min, and eluted in 30 µL of 0.1x TE. Final libraries were quantified and assessed for quality using a High Sensitivity D5000 Screen Tape Assay run on the 4150 TapeStation system (Agilent Technologies, Santa Clara, United States).

### Library preparation for Multi-Read&Tag using primer-timed signal augmentation

Primer-timed signal augmentation (PTSA) was implemented by sequentially adding primers spanning the P5-Forward or P7-Reverse sequences and reader-specific barcodes during enrichment PCR. The order and timing of primer addition were determined in advance using quantitative real-time PCR (qPCR). For qPCR optimization, 21 µL of purified multiplex-tagmentation DNA was combined with 2 µL of P5-Forward and P7-Reverse primers (10 µM concentration) and 25 µL of NEBNext HiFi 2x PCR Master Mix. The reactions were then subjected to a preamplification PCR with the following program: Incubation at 72 °C for 5 min (gap-filling) and 98 °C for 30 sec; then 5-8 cycles of 98 °C for 10 sec, 55 °C for 10 sec and 63 °C for 45 sec; and a final extension at 72 °C for 5 min, with eventual cooling to 4 °C. DNA was purified by addition of 65 µL SPRI paramagnetic beads as described before, and subsequently used for qPCR. For each reader in the multiplexed reaction, three technical qPCR replicates were performed. qPCR reactions contained 5 µL Fast SYBR Green Master Mix (applied biosystems by Thermo Fisher Scientific, Cat.No. 4385612), 1 µL of reader-specific primer mix (5 µM concentration), and 2 µL pre-amplified DNA. The qPCR program was started with 95 °C for 30 sec, followed by 35 cycles of 98 °C for 15 sec and 68 °C for 75 sec. The resulting Cp values were used to calculate the PTSA schedule (for more details, see Step 72 of the protocols.io version of the Read&Tag protocol).

For final Multi-Read&Tag library amplification, reactions were set up using primers specific to the lowest-abundant histone modification. 21 µL of purified multiplex-tagmentation DNA was combined with 0.8 µL of reader-specific forward and reverse oligonucleotides each, followed by addition of 25 µL NEBNext HiFi 2x PCR Master Mix. The enrichment PCR was carried out with an initial gap-filling step (72 °C for 5 min), followed by denaturation at 98 °C for 30 sec and a number of PTSA-scheduled amplification cycles (98 °C for 10 sec, 72 °C for 75 sec), ending with a final extension at 72 °C for 5 min. During amplification, cycling was paused at the end of the 72 °C step to add 0.8 µL of reader-specific forward/reverse primers for higher-abundance modifications according to the PTSA schedule. The exact cycle numbers used for each target in this publication are provided in Supplementary Table S5. The obtained libraries were purified by a size selection approach to remove large fragments. For this, 20 µL of SPRI paramagnetic beads were added to the PCR reaction and incubated for 7 min. After magnetic separation, the supernatant was transferred into a fresh reaction tube and mixed with 45 µL of fresh SPRI paramagnetic beads, again incubated for 7 min at RT, washed twice with 80 % ethanol, air-dried for 2 min, and eluted in 25-30 µL of 0.1x TE. Final libraries were quantified and assessed for quality using a High Sensitivity D5000 Screen Tape Assay run on the 4150 TapeStation system (Agilent Technologies, Santa Clara, United States).

### Mouse liver nuclei enrichment

Livers were enriched from male C57BL/6N mice (8-10 weeks old). Mice were housed under specific-pathogen-free conditions in individually ventilated cages with a 12-hour light/dark cycle, and were provided with sterile food and water ad libitum. All procedures were approved by the local government authorities and conducted in accordance with the European law on the protection of animal welfare (animal facility registered for breeding and the use of animals for scientific purpose KVR-I/221-TA166/22_03-06). Following whole-tissue dissection, livers were snap-frozen in liquid nitrogen and kept at -80°C until use. For nuclear enrichment, the FrozONE workflow ^42^ was used. In detail, frozen livers were dissected over dry ice into 150 mg pieces and transferred into 2 ml tubes. One stainless steel ball and 1 ml cold Nuclei EZ lysis buffer (Sigma-Aldrich, Cat.No. NUC101) supplemented with PIC were added. Immediately after the addition of lysis buffer, livers were homogenized using the QIAGEN TissueLyser II at 30 Hz for 15 sec. Homogenates were transferred to 50 mL Falcon tubes with a 40 µm cell strainer and centrifuged for 2 min at 100 x g. Flow-throughs were transferred to 1.5 mL tubes on ice and incubated for 5 min. Lysates were then centrifuged at 500 x g at 4 °C for 5 min. Supernatants were carefully removed and discarded. Nuclei pellets were dissociated by moving tubes across a rack 10 times and were resuspended by adding 1 mL of cold Nuclei EZ lysis buffer and inverting 5 times. From this step, the cycle of ice incubation, centrifugation, aspiration, and resuspension was repeated a total of three more times. The final resuspension was done in Read&Tag wash buffer. Subsequently, Read&Tag was performed as described above.

### Chromatin immunoprecipitation (ChIP)

HA36CB1 mESCs (20-30 x 10⁶) were trypsinized using TrypLE (Gibco, Cat.No. 12605-028), washed once in 1x PBS, and fixed with 1 % formaldehyde for 8 min at RT, followed by reaction quenching with the addition of glycine to a final concentration of 0.12 M and incubation on ice for 10 min. Cells were washed twice in 10 mL of ice cold 1x PBS and then centrifuged at 680 x g for 5 min. Cells were resuspended and incubated for 10 min in 10 mL Paro Rinse 1 buffer (10 mM Tris pH 8.0, 10 mM EDTA, 0.5 mM EGTA) on ice, then centrifuged at 680 x g for 5 min. Cells were resuspended in 10 mL of Paro Rinse 2 buffer (10 mM Tris pH 8.0, 0.25 % Triton X-100, 1 mM EDTA, 0.5 mM EGTA, and 200 mM NaCl), filtered three times through a prechilled G26 syringe, and incubated on ice for 10 min, followed by centrifugation at 680 x g for 5 min. Final cell lysis was performed with Mnase Digest buffer (20 mM Tris pH 8.0, 5 mM MgCl_2_, 1 mM CaCl_2_, 10 mM NaCl, 0.25 M Sucrose, 1 % Triton X-100, supplemented with 1x PIC at a concentration-dependent volume (20,000 cells per µL) for 30 min on ice. For MNase digestion, samples were incubated in a water bath at 37 °C for 2 min. 50 U MNase (Sigma Aldrich, Cat.No. N5386) per 1 mL of buffer was added, and the samples were incubated for a further 20 min at 37 °C in a water bath. During incubation, the reaction tubes were inverted 2 times in 5 min intervals. Digestion was stopped with NaCl, EDTA, EGTA, and SDS (final: 500 mM NaCl, 20 mM EDTA, 10 mM EGTA, 0.1 % SDS). Chromatin was further sonicated using a Diagenode Bioruptor Pico device (4 cycles, 32 sec ON/50 sec OFF) and clarified by centrifugation at 12,000 x g for 10 min; the supernatant was then used for immunoprecipitation.

For each ChIP, 100 µg chromatin was incubated with 30 µl Protein A+G magnetic beads (Sigma Aldrich, Magna ChIP, Cat.No. 16-663). Beads were washed twice with 1x TE buffer (10 mM Tris-HCl pH 8.0, 1 mM EDTA) and blocked for 2 h in 1x TE buffer supplemented with recombinant Albumin (20 µg/mL; New England Biolabs, Cat.No. B9200S), denatured yeast tRNA (1 mg/mL; heated at 95 °C for 5 min; Thermo Fisher Scientific, Cat.No. 15401011), and 1x PIC. Beads were washed three times with 1x TE buffer, twice with MNase digestion buffer, and finally resuspended in 1x TE buffer at the original bead volume.

Chromatin was pre-cleared with 10 µL blocked Protein A+G beads for 1 h with overhead rotation at 4 °C. 100 µg of pre-cleared chromatin was transferred to fresh tubes and incubated overnight at 4 °C with 5 µg of either anti-H3K9me3 (Abcam, Cat.No. ab8898), anti-H3K27me3 (Diagenode, Cat.No. C15410195), or anti-H3K4me3 (Abcam, Cat.No. ab8580) antibody. The remaining 20 µL blocked beads were added the next day, and samples were incubated for an additional 3 h with overhead rotation at 4 °C. Beads were washed twice at RT with a high-salt buffer (50 mM HEPES-KOH pH 7.5, 500 mM NaCl, 1 mM EDTA, 1 % Triton X-100, 0.1 % sodium deoxycholate, 0.1 % SDS), once with DOC buffer (250 mM LiCl, 0.5 % NP-40 (IGEPAL CA-630, Sigma Aldrich, Cat.No. I8896-50ML), 0.5 % sodium deoxycholate, 1 mM EDTA, 10 mM Tris-HCl pH 8.0), and twice with 1x TE buffer. Beads were transferred to fresh tubes before the final wash and resuspended in 50 µL 1x TE buffer. Samples were treated with 10 µg RNase A (Roche, Cat.No. 10109142001) for 30 min at 37 °C with shaking at 1100 rpm, then incubated with 50 µg Proteinase K and SDS (final concentration 0.54 %) for 2 h at 68 °C and 1100 rpm. Beads were separated magnetically, and DNA was purified from the supernatant using 100 µl SPRI paramagnetic beads (AMPure XP beads by Beckman Coulter; alternatively, MAGFLO NGS beads by Integra) following the manufacturer’s instructions. DNA was eluted in 50 µl of 0.1x TE buffer.

### ChIP-seq library preparation

ChIP-seq libraries were prepared using the NEBNext Ultra II DNA Library Prep Kit for Illumina (New England Biolabs, Cat.No. E7645) following the manufacturer’s instructions. Libraries were analyzed using a High Sensitivity D5000 Screen Tape Assay run on the 4150 TapeStation system (Agilent Technologies, Santa Clara, United States).

### Chromatin accessibility profiling by ATAC-seq

ATAC-seq was performed on HA36CB1 cells as previously described ^63^, with minor modifications. Cells were dissociated, counted, and 50,000 cells per sample were pelleted. Cell pellets were resuspended and lysed in 50 µL of cold ATAC-seq resuspension buffer 1 (ATAC-RSB1; 10 mM Tris-HCl, pH 7.5, 10 mM NaCl, 3 mM MgCl₂, 0.1% NP-40, 0.1% Tween-20 [Sigma Aldrich, Cat.No. P1379-25ML], 0.01% digitonin) for 3 minutes. Subsequently, 1 mL of cold ATAC-seq resuspension buffer 2 (ATAC-RSB2; 10 mM Tris-HCl, pH 7.5, 10 mM NaCl, 3 mM MgCl₂, 0.1% Tween-20) was added to dilute ATAC-RSB1. Nuclei were pelleted by centrifugation at 500 g for 10 minutes at 4°C and resuspended in 1x transposition mix (Illumina; 25 µL Tagment DNA Buffer, 2.5 µL Tagment DNA Enzyme, 16.5 µL PBS, 0.25 µL 2% digitonin, 0.5 µL 10% Tween-20, 5.25 µL nuclease-free water). The transposition reaction was incubated for 30 minutes at 37°C with shaking at 1,000 rpm. DNA was immediately purified using the Qiagen MinElute PCR Purification Kit (Cat.No. 28004) according to the manufacturer’s instructions and eluted in 10 µL of 0.1x TE buffer. Libraries were amplified using sequencing primers and NEBNext Q5 Hot Start HiFi 2x PCR Master Mix in a 50 µL reaction for an initial 4 cycles. To determine the optimal number of amplification cycles, 4 µL of the reaction mixture was removed for qPCR analysis, and the remaining 46 µL was amplified. The amplified libraries were purified using the Qiagen PCR Purification Kit and eluted in 20 µL of 0.1x TE buffer. Size selection was performed using AMPure XP beads to enrich for nucleosome-free and mono-nucleosome fragments. Samples were first purified with 0.6x AMPure XP beads, and the unbound fraction was subsequently purified with 2.68x AMPure XP beads. Final libraries were eluted, quantified, and assessed for quality using a High Sensitivity D5000 Screen Tape Assay run on the 4150 TapeStation system.

### PolyA RNA-sequencing (mRNA-seq)

Total RNA was isolated from HA36CB1 cells using the RNeasy Plus mini kit (Qiagen, Cat.No. 74134). RNA integrity was measured using a 4150 TapeStation system. PolyA-tailed mRNAs were isolated and enriched from 0.5 µg of total RNA using the NEBNext Poly(A) mRNA Magnetic Isolation Module (New England Biolabs, Cat.No. E7490), according to the manufacturer’s instructions. Libraries were prepared using the NEBNext Ultra II Directional RNA Library Prep Kit for Illumina (New England Biolabs, Cat.No. E7760), following the manufacturer’s instructions. Final libraries were quantified and assessed for quality using the 4150 TapeStation system.

### Next-generation sequencing

Libraries with unique index barcodes were combined in equal molar ratios and sequenced as pools on an Illumina NextSeq 1000 or NextSeq 2000 in 60 base-pair paired-end mode at the Laboratory of Functional Genomic Analysis (LAFUGA, Gene Center Munich, LMU). Library demultiplexing was performed following Illumina standards.

### Genomics data analysis

#### Short read processing

We used the snakePipes (v2.7.0) automated, reproducible workflows, built with Snakemake and Python, for the preprocessing of NGS datasets ^64^. Short reads and publicly available datasets were mapped to the mouse genome (GRCm38; mm10) using Bowtie2 (v2.3.3.1) ^65^. The DNAmapping pipeline was additionally set to: ‘--mapq 3’ and ‘--dedup’. For the CUT&Tag and Read&Tag datasets, the ‘--cutntag’ flag was used for mapping with parameters used in Kaya-Okur et al. 2019: ’--local --very-sensitive-local --no-mixed --no-discordant --phred33 -I 10 -X 700’. Reads were sorted using SAMtools (v1.9) ^66^ and duplicates were removed by picard tools (v2.9.2) ^67^. Coverage bigWig files were generated using bamCoverage, CPM-normalized using ‘--normalizeUsing CPM’, and CPM-normalized BigWig files were averaged using bigwigAverage from deepTools (v3.5.6) ^68^.

### Integrative genome viewer (IGV) snapshots

CPM-normalized BigWig files were loaded into IGV (https://igv.org/app/#) and visualized as group auto-scaled bars when the same assay was used to target a specific chromatin mark. Genome snapshots were saved as SVG files for further editing with Affinity Designer 2 software (v2.1.0).

### Peak calling

Peak calling for ChIP-seq datasets was performed with the ChIPseq pipeline from snakePipes (v2.7.0) and MACS2 (v2.1.2) ^35^ using input samples as control and the flags ‘broad: True’ and ‘--peakCallerOptions ’--qvalue 0.00001’’. For CUT&Tag and Read&Tag peak calling with snakePipes/MACS2, the parameters used were: macs2 callpeak -t input_file -p 1e-5 -f BEDPE/BED (Paired End vs. Single End sequencing data) -keep-dup all -n out_name. For CUT&Tag and Read&Tag peak calling with SEACR (v1.3) ^36^, the parameters used were: bash SEACR_1.3.sh target.bedgraph control.bedgraph norm stringent output. For CUT&Tag and Read&Tag narrow peak calling with GoPeaks ^37^, we used the flag ’--pval 0.001’. For CUT&TAG and Read&Tag broad peaks calling with GoPeaks, we used the parameters: ’--broad --mdist 3000 --pval 0.001’.

### Genome-wide binning and signal quantification

To enable genome-wide comparisons among Read&Tag, ChIP-seq, and CUT&Tag datasets, the mouse reference genome (mm10) was partitioned into non-overlapping 1-kb intervals across all standard autosomes, excluding sex chromosomes, using *BSgenome.Mmusculus.UCSC.mm10 (1.4.3)* ^69^. Windows were generated across a predefined set of chromosomes and represented as *GRanges* objects. Chromosome naming was standardized to UCSC style. For each 1-kb window, we additionally computed several genomic covariates, including mappability, blacklist overlap, whole-genome bisulfite sequencing (WGBS)-derived CpG methylation statistics, GC fraction and CpG density. Mappability was calculated from a BED track containing per-base or interval-level mappability scores by computing a width-weighted mean score across each 1-kb window, with uncovered bases treated as zero. Blacklist regions were imported from a BED file and encoded as a binary flag indicating whether a given window overlapped a blacklist interval. Sequence covariates were computed directly from the mm10 reference genome: GC fraction was defined as the fraction of G and C bases per 1-kb window, and CpG density as the number of “CG” dinucleotides per base pair in that window. WGBS-derived methylation features were obtained from bigWig signal tracks (see Data availability) and summarized per 1-kb window as the mean methylation beta value across CpG loci, the number of CpG loci, and the estimated number of methylated CpG sites above a predefined methylation threshold. Where indicated, windows overlapping blacklist regions were masked by setting feature values to missing. Signal tracks in bigWig format were summarized over the same 1-kb windows using *rtracklayer (1.68.0)* ^70^. For each track, the signal was imported chromosome by chromosome over the full genomic span covered by the corresponding window set, and overlap between signal intervals and windows was calculated using *GenomicRanges (1.60.0)* ^71^. For each window, we computed either the base-pair-weighted mean signal or the integrated signal, depending on the downstream application. Unless otherwise specified, the reported signal corresponds to the base-pair-weighted mean signal per 1-kb window. When bigWig files contained only explicit non-zero intervals, missing positions were treated as implicit zeros by default, consistent with deepTools (v3.5.6) ^68^ bamCoverage outputs. Optionally, windows with insufficient covered fraction in a given bigWig track were masked using a minimum coverage-fraction threshold. Windows overlapping blacklist regions were also masked during signal extraction when specified.

For genome-wide correlation analyses, summarized 1-kb signals were transformed using a log2(x + 1) transformation to stabilize variance and reduce the influence of highly enriched intervals. Pairwise Pearson and Spearman correlation coefficients were then calculated across matched 1-kb windows between biological replicates, between reader constructs, and between Read&Tag and matched reference datasets. Depending on the figure, correlations were visualized either as heatmaps or as scatter-density plots. All genome-wide feature generation, bigWig summarization, and correlation analyses were implemented in R using Bioconductor packages, including *GenomicRanges (1.60.0)*, *GenomeInfoDb (1.44.3)* ^72^, *rtracklayer (1.68.0)*, *BSgenome.Mmusculus.UCSC.mm10 (1.4.3)* and *Biostrings (2.76.0)* ^73^. Correlation matrices were visualized as heatmaps or scatter plots using R packages.

### Fraction of reads in peaks (FRiP)

FRiP scores were calculated to assess target-specific signal enrichment for Read&Tag, CUT&Tag, and ChIP-seq datasets. FRiP was defined as the fraction of sequenced reads or fragments overlapping a predefined set of peak regions for the corresponding chromatin feature. For each chromatin mark, peak regions were merged across replicates and reduced to non-overlapping intervals prior to FRiP calculation to avoid double-counting of overlapping features. Only peaks located on standard chromosomes were retained for downstream analysis. FRiP analysis was performed directly on coordinate-sorted BAM files using the R packages *Rsamtools (2.24.1)* ^74^, *GenomicAlignments (1.44.0)* ^75^, and *GenomicRanges (1.60.0)* ^71^. Reads were filtered to remove unmapped, secondary, and supplementary alignments. Only reads with mapping quality ≥3 were retained. Duplicate reads were either retained or excluded depending on the analysis, as indicated in the figure legends. For paired-end datasets, FRiP was computed at the fragment level. Properly paired reads were required, and fragments were defined using inferred template lengths. For single-end datasets, FRiP was computed at the read level, with reads extended to a fixed length of 150 bp in the 3′ direction to approximate fragment coverage. Filtered reads or fragments were converted to genomic intervals and intersected with the corresponding peak sets. A read or fragment was considered to overlap a peak if it intersected at least one peak interval by ≥1 bp. FRiP was calculated as:

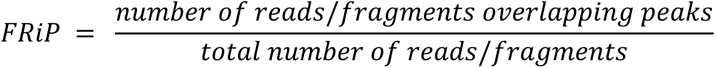

FRiP scores were computed separately for each sample, peak set, and duplicate-handling mode. For analyses comparing duplicate-retained and duplicate-removed data, FRiP was reported for both conditions.

### Precision-recall analysis of peak recovery

To benchmark peak recovery across peak-calling strategies and chromatin marks, we performed peak-level precision-recall analysis for H3K4me3, H3K9me3, and H3K27me3. For each mark, a fixed reference peak set was defined from matched ChIP-seq data using MACS2 and replicate-consensus peak merging (blacklist-filtered). Test peak sets were generated from Read&Tag datasets using GoPeaks, SEACR, or MACS2 run with CUT&Tag-recommended parameters (see the “Peak calling” methods section for details). Each test peak set was compared only to the reference peak set for the corresponding chromatin mark. Precision and recall were computed using binary peak overlaps, without imposing a minimum reciprocal overlap fraction, to quantify peak recovery independently of peak width. For a given test peak set *T* and reference peak set *R*, overlaps were defined as follows:

A test peak was counted as “recovered” if it overlapped at least one reference peak (≥100 bp). A reference peak was counted as “recalled” if it overlapped at least one test peak (≥100 bp). Overlaps were calculated using GenomicRanges::countOverlaps with type = "any". For each test peak set, precision and recall were computed at the peak level as:

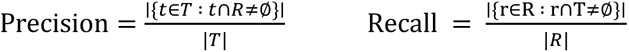

Metrics were reported as percentages and computed independently for each chromatin mark and each test peak set.

### Peak-width and peak-count diagnostics

Because peak-level overlap metrics can be influenced by peak-caller behavior for broad chromatin domains, we also quantified peak-width distributions for all reference and test peak sets. For each peak set, peak widths were calculated from interval lengths and summarized as median width and number of peaks. These diagnostics helped to interpret precision-recall metrics, especially for broad marks (H3K27me3 and H3K9me3), for which over-fragmentation or overcalling can inflate recall while reducing precision.

### Consensus peaks

Peaks were imported *with rtracklayer (1.68.0)* and handled as GRanges objects *(GenomicRanges 1.60.0)*, restricted to autosomes and trimmed against the corresponding ENCODE blacklist using GenomicRanges::setdiff before and after merging. A peak was counted as supported by another replicate when at least 25% of its width was covered by peaks of that replicate and it overlapped at least one peak of that replicate whose own width was at least 25% reciprocally covered. Overlapping base pairs were summed per peak before thresholding, and support was evaluated independently for each pair of replicates, making the result invariant to replicate order. Regions supported by at least *k* of *n* replicates were retained as the merged union of the supporting peaks (*GenomicRanges::reduce*, min.gapwidth = 1). Regions shorter than 150 bp (H3K4me3, Bivalent) or 450 bp (H3K27me3, H3K9me3) were discarded. For ChIP-seq, MACS2 peaks with a score of at least 10 were required in both replicates (*k* = *n* = 2). For Read&Tag, peaks were required in all 3 replicates (*k* = *n* = 3) for each mark and each caller, and consensus regions were obtained by combining the GoPeaks and SEACR sets by range reduction. Published reChIP-seq libraries (GSE242686)^52^ were reanalyzed with the same rule, requiring support from both replicates within each antibody order and then from both orders. This reproducibility-filtered reanalysis is more conservative than the published peak set.

### Spike-in normalization

Reads were aligned with Bowtie2 (v2.3.3.1) to a concatenated reference comprising the mouse (mm10) and *Drosophila melanogaster* (dm6) genomes, in which spike-in contigs were given a distinguishing suffix. After filtering/deduplication as described above, endogenous and spike-in read counts were obtained per sample from the BAM index using samtools idxstats (v1.9). For each sample *i*, the ratio of endogenous to spike-in reads Rᵢ = Nᵢ/Sᵢ was computed, and a scale factor defined as sᵢ = 10⁶/(Sᵢ × G), where *G* is the geometric mean of *R* across the 0 h untreated samples, so that 0 h signal is placed at CPM-equivalent levels and all other samples are expressed relative to it. Division by *G* places the normalized signal on a scale numerically comparable to counts per million (CPM) while preserving the relative differences reported by the spike-in. Where the number of spike-in nuclei differed systematically between processing batches, *G* was determined independently within each batch from the 0 h sample of that batch, removing a constant multiplicative offset between batches while preserving the condition differences measured within each batch. Normalized coverage tracks were generated with deepTools bamCoverage (v3.5.0) using 10-bp bins, --normalizeUsing None and --scaleFactor set to sᵢ, ensuring the spike-in factor was the only scaling applied.

### Statistics and reproducibility

The number of independent biological replicates (n) for each dataset is given in the figure legends and Supplementary Table S8. Read&Tag with the monovalent readers, their binding-deficient mutants, the combinatorial readers and the RNAP2-Ser2ph reader was performed with two to four biological replicates in cells or tissue. Multi-Read&Tag, low cellular input Read&Tag, and the PRC1-degron time course were performed with two biological replicates. Reference datasets (RNA-seq, ChIP-seq, CUT&Tag and MeDIP) comprise at least two biological replicates (except for liver H3K4me3 ChIP-seq [n=1]). ATAC-seq (n = 1) was used only to define accessible chromatin regions. For cell line experiments, biological replicates represent independently cultured cell populations processed on separate days. For tissue experiments, biological replicates correspond to independent tissue samples derived from distinct animals. Unless otherwise specified, no technical replicates were averaged prior to analysis.

Genome-wide analyses were performed on fixed 1-kb windows or predefined peak sets. Windows within a library are not independent observations. They are units of measurement from which one summary value per library is derived, such as a Pearson or Spearman correlation coefficient, the fraction of reads in peaks (FRiP), or precision and recall against a reference peak set. The biological replicate is therefore the unit of reproducibility. Summary metrics were computed for each replicate separately and are shown as individual replicate values together with the mean and the range (minimum to maximum). Some intervals were derived in other ways: by resampling genomic regions within a library (Supplementary Table S6), or from pairings of replicates that share libraries (Supplementary Fig. 1f). These are described as such and are not presented as confidence intervals for biological variation. The aim of this study was to assess the reproducibility of Read&Tag and its concordance with established assays. For this reason, and given the number of replicates, results are reported descriptively, and no formal hypothesis tests or multiple-testing corrections were performed. No assumption of normality was made for genome-wide distributions, and non-parametric summaries were used where appropriate. Aggregate profiles show the mean signal across regions at each position (averaged across replicates, as indicated in the figure legends). Unless stated otherwise, summary statistics include all windows or regions that passed the quality filters described above (mappability, blacklist and chromosome filters). All samples were processed with identical preprocessing, filtering and normalization.

## Data availability

Publicly available ChIP-seq, CUT&Tag, ATAC-seq, MeDIP and RNA-seq datasets used in this study, from mouse ES cells, mouse liver and human MCF7 cells, are listed in Supplementary Table S7. The Read&Tag, ChIP-seq, ATAC-seq, and RNA-seq data generated in this study are listed in Supplementary Table S8 and are available at ArrayExpress under accession E-MTAB-16952, with public release on 2027-05-04.

BigWig of CpG beta values from WGBS data in mESCs (129xC57BL/6) ^76^ was obtained from the DNA methylation hub maintained by the Smith lab (https://hgdownload.soe.ucsc.edu/hubs/methbase/v1/mm10/Mouse_ESC.meth.bw). Blacklist regions for filtering were obtained from the Boyle lab (https://github.com/Boyle-Lab/Blacklist/blob/master/lists/mm10-blacklist.v2.bed.gz) ^77^. Mouse mm10 Umap single-read mappability track for 100 bp ^78^ was downloaded from here: https://bismap.hoffmanlab.org/. We downloaded reference peak sets from 2-month-old mouse livers from the ENCODE portal ^79^ (https://www.encodeproject.org/) with the following identifiers: ENCFF420PUU (H3K4me3), ENCFF251NKN (H3K9me3), and ENCFF777YCC (H3K27me3). Uncropped and unprocessed scans of all Western blots are provided in Source Data.

## Code availability

R code has been deposited on Zenodo (https://doi.org/10.5281/zenodo.23059645).

## Supporting information

Schneider_et_al_supplementary_figures_biorxiv_2026

Schneider_et_al_supplementary_tables_biorxiv_2026

## Acknowledgements

We thank Peter Becker, Philipp Korber, Tamas Schauer, Michela Palamin, Colm Doyle, and Arnaud Krebs for their comments on the manuscript. We thank all current and former members of the Villaseñor lab and the Department of Molecular Biology for their discussions and insights on the project. We thank Tobias Straub from the BMC Bioinformatics Unit for providing access to and maintaining the computational cluster. We thank Stefan Krebs and the LMU LAFUGA team for their assistance with Illumina sequencing. This work was supported by the Deutsche Forschungsgemeinschaft (DFG, German Research Foundation) [468577222 to R.V.], the Friedrich-Baur-Stiftung [Project number 21499], the LMU Munich’s Institutional Strategy LMU excellent within the framework of the German Excellence Initiative, and by the European Research Council [EpiCblood, 101115861 to R.V.]. Views and opinions expressed are however those of the author(s) only and do not necessarily reflect those of the European Union or the European Research Council Executive Agency. Neither the European Union nor the granting authority can be held responsible for them. M.S.R. is also supported by the Deutsche Forschungsgemeinschaft [213249687 to M.S.R.], SFB 1064 “Chromatin Dynamics”, INST 86/1800-1 FUGG, and LMU Munich’s Institutional Strategy LMU excellent within the framework of the German Excellence Initiative.

## Author contributions

M.S. developed materials and the Read&Tag protocol. M.S., L.C.K., C.S., N.R.B., C.B., and A.C.S. performed experiments. L.A.H. and M.S.R. provided mouse liver samples and protocols for nuclei enrichment from frozen tissue. R.V., M.S., and L.C.K. analyzed the data. L.C.K. wrote the protocols.io Read&Tag protocol with input from all authors. R.V. designed, supervised the study, and wrote the manuscript with input from all authors. All authors discussed the results and commented on the manuscript.

## Corresponding author

Correspondence and requests for materials should be addressed to R.V.

## Data access

The genome browser tracks shown in **Figure 1** of this study can be accessed here.

The genome browser tracks shown in **Figure 2** of this study can be accessed here.

The genome browser tracks shown in **Figure 3** of this study can be accessed here.

The genome browser tracks shown in **Figure 4** of this study can be accessed here.

The genome browser tracks shown in **Figure 5** of this study can be accessed here.

The genome browser tracks shown in **Figure 6** of this study can be accessed here.

The genome browser tracks shown in **Figure 7** of this study can be accessed here.

## References

1. Henikoff, S. & Ahmad, K. In situ tools for chromatin structural epigenomics. Protein Sci. 31, e4458 (2022).

2. Skene, P. J. & Henikoff, S. An efficient targeted nuclease strategy for high-resolution mapping of DNA binding sites. eLife 6, e21856 (2017).

3. Harada, A. et al. A chromatin integration labelling method enables epigenomic profiling with lower input. Nat. Cell Biol. 21, 287–296 (2019).

4. Kaya-Okur, H. S. et al. CUT&Tag for efficient epigenomic profiling of small samples and single cells. Nat. Commun. 10, 1930 (2019).

5. Carter, B. et al. Mapping histone modifications in low cell number and single cells using antibody-guided chromatin tagmentation (ACT-seq). Nat. Commun. 10, 3747 (2019).

6. Wang, Q. et al. CoBATCH for High-Throughput Single-Cell Epigenomic Profiling. Mol. Cell 76, 206–216.e7 (2019).

7. Gopalan, S., Wang, Y., Harper, N. W., Garber, M. & Fazzio, T. G. Simultaneous profiling of multiple chromatin proteins in the same cells. Mol. Cell 81, 4736–4746.e5 (2021).

8. Carter, B., Ku, W. L., Pelt, J. & Zhao, K. Concurrent mapping of multiple epigenetic marks and co-occupancy using ACT2-seq. Cell Biosci. 11, 198 (2021).

9. Xiong, H., Wang, Q., Li, C. C. & He, A. Single-cell joint profiling of multiple epigenetic proteins and gene transcription. Sci. Adv. 10, eadi3664 (2024).

10. Meers, M. P., Llagas, G., Janssens, D. H., Codomo, C. A. & Henikoff, S. Multifactorial profiling of epigenetic landscapes at single-cell resolution using MulTI-Tag. Nat. Biotechnol. 41, 708–716 (2023).

11. Stuart, T. et al. Nanobody-tethered transposition enables multifactorial chromatin profiling at single-cell resolution. Nat. Biotechnol. 41, 806–812 (2023).

12. Bartosovic, M. & Castelo-Branco, G. Multimodal chromatin profiling using nanobody-based single-cell CUT&Tag. Nat. Biotechnol. 41, 794–805 (2023).

13. Dimitriu, M. A., Arzate-Mejía, R. G., Steg, L. C., Germain, P.-L. & Mansuy, I. M. NanoTag is an improved method to map interactions between DNA and proteins not requiring IgG. Sci. Rep. 15, 28672 (2025).

14. Wang, X. et al. Af-CUT&Tag: a sensitive and antibody-free chromatin profiling method using genetically encoded tags and high-affinity binders fused to Tn5. Nat. Commun. 10.1038/s41467-026-68454-9 (2026) doi:10.1038/s41467-026-68454-9.

15. Park, P. J. ChIP–seq: advantages and challenges of a maturing technology. Nat. Rev. Genet. 10, 669–680 (2009).

16. Nishikori, S. et al. Broad Ranges of Affinity and Specificity of Anti-Histone Antibodies Revealed by a Quantitative Peptide Immunoprecipitation Assay. J. Mol. Biol. 424, 391– 399 (2012).

17. Egelhofer, T. A. et al. An assessment of histone-modification antibody quality. Nat. Struct. Mol. Biol. 18, 91–93 (2011).

18. 18. Wang, M. & Zhang, Y. Tn5 transposase-based epigenomic profiling methods are prone to open chromatin bias. Preprint at 10.1101/2021.07.09.451758 (2021).

19. Musselman, C. A., Lalonde, M.-E., Côté, J. & Kutateladze, T. G. Perceiving the epigenetic landscape through histone readers. Nat. Struct. Mol. Biol. 19, 1218–1227 (2012).

20. Moore, K. E. et al. A General Molecular Affinity Strategy for Global Detection and Proteomic Analysis of Lysine Methylation. Mol. Cell 50, 444–456 (2013).

21. Su, Z. et al. ChIP-less analysis of chromatin states. Epigenetics Chromatin 7, 7 (2014).

22. Mauser, R., Kungulovski, G., Keup, C., Reinhardt, R. & Jeltsch, A. Application of dual reading domains as novel reagents in chromatin biology reveals a new H3K9me3 and H3K36me2/3 bivalent chromatin state. Epigenetics Chromatin 10, 45 (2017).

23. Delachat, A. M.-F. et al. Engineered Multivalent Sensors to Detect Coexisting Histone Modifications in Living Stem Cells. Cell Chem. Biol. 25, 51–56.e6 (2018).

24. Tekel, S. J. et al. Tandem Histone-Binding Domains Enhance the Activity of a Synthetic Chromatin Effector. ACS Synth. Biol. 7, 842–852 (2018).

25. Albanese, K. I. et al. Engineered Reader Proteins for Enhanced Detection of Methylated Lysine on Histones. ACS Chem. Biol. 15, 103–111 (2020).

26. Villaseñor, R. et al. ChromID identifies the protein interactome at chromatin marks. Nat. Biotechnol. 38, 728–736 (2020).

27. Veggiani, G. et al. High-affinity chromodomains engineered for improved detection of histone methylation and enhanced CRISPR-based gene repression. Nat. Commun. 13, 6975 (2022).

28. Zhao, H. et al. Manipulating Cation−π Interactions of Reader Proteins in Living Cells with Genetic Code Expansion. J. Am. Chem. Soc. 145, 16406–16416 (2023).

29. Chong, S. et al. Single-column purification of free recombinant proteins using a self-cleavable affinity tag derived from a protein splicing element. Gene 192, 271–281 (1997).

30. Vermeulen, M. et al. Selective Anchoring of TFIID to Nucleosomes by Trimethylation of Histone H3 Lysine 4. Cell 131, 58–69 (2007).

31. Jacobs, S. A. & Khorasanizadeh, S. Structure of HP1 Chromodomain Bound to a Lysine 9-Methylated Histone H3 Tail. Science 295, 2080–2083 (2002).

32. Li, R., Grimm, S. A. & Wade, P. A. CUT&Tag-BS for simultaneous profiling of histone modification and DNA methylation with high efficiency and low cost. *Cell Rep*. Methods 1, 100118 (2021).

33. Gjaltema, R. A. F. et al. Distal and proximal cis-regulatory elements sense X chromosome dosage and developmental state at the Xist locus. Mol. Cell 82, 190–208.e17 (2022).

34. Park, B. J. et al. CUT&Tag overcomes biases of ChIP and establishes chromatin patterns for repetitive genomic loci. iScience 28, 113757 (2025).

35. Zhang, Y. et al. Model-based Analysis of ChIP-Seq (MACS). Genome Biol. 9, R137 (2008).

36. Meers, M. P., Tenenbaum, D. & Henikoff, S. Peak calling by Sparse Enrichment Analysis for CUT&RUN chromatin profiling. Epigenetics Chromatin 12, 42 (2019).

37. Yashar, W. M. et al. GoPeaks: histone modification peak calling for CUT&Tag. Genome Biol. 23, 144 (2022).

38. Dobrinić, P., Szczurek, A. T. & Klose, R. J. PRC1 drives Polycomb-mediated gene repression by controlling transcription initiation and burst frequency. Nat. Struct. Mol. Biol. 28, 811–824 (2021).

39. The ENCODE Project Consortium. An integrated encyclopedia of DNA elements in the human genome. Nature 489, 57–74 (2012).

40. Tian, C. et al. Impaired histone inheritance promotes tumor progression. Nat. Commun. 14, 3429 (2023).

41. Shi, Y., Gong, W., Gong, X., Wang, P. & Zhao, X. Genome-wide DNA methylation analysis of breast cancer MCF-7 / Taxol cells with MeDIP-Seq. PLOS ONE 15, e0241515 (2020).

42. Huschet, L. A. et al. FrozONE: quick cell nucleus enrichment for comprehensive proteomics analysis of frozen tissues. Life Sci. Alliance 8, e202403130 (2025).

43. Wang, S. et al. Epigenetic Compensation Promotes Liver Regeneration. Dev. Cell 50, 43–56.e6 (2019).

44. Yang, N. et al. A hyper-quiescent chromatin state formed during aging is reversed by regeneration. Mol. Cell 83, 1659–1676.e11 (2023).

45. Uchino, S. et al. Live imaging of transcription sites using an elongating RNA polymerase II–specific probe. J. Cell Biol. 221, e202104134 (2022).

46. Guo, Y. E. et al. Pol II phosphorylation regulates a switch between transcriptional and splicing condensates. Nature 572, 543–548 (2019).

47. Abe, K., Schauer, T. & Torres-Padilla, M.-E. Distinct patterns of RNA polymerase II and transcriptional elongation characterize mammalian genome activation. Cell Rep. 41, 111865 (2022).

48. Padrão, N., et al. TRIM24 as a therapeutic target in endocrine treatment–resistant breast cancer. Proc. Natl. Acad. Sci. 122, e2507571122 (2025).

49. Bernstein, B. E. et al. A Bivalent Chromatin Structure Marks Key Developmental Genes in Embryonic Stem Cells. Cell 125, 315–326 (2006).

50. Mikkelsen, T. S. et al. Genome-wide maps of chromatin state in pluripotent and lineage-committed cells. Nature 448, 553–560 (2007).

51. Macrae, T. A., Fothergill-Robinson, J. & Ramalho-Santos, M. Regulation, functions and transmission of bivalent chromatin during mammalian development. Nat. Rev. Mol. Cell Biol. 24, 6–26 (2023).

52. Seneviratne, J. A., Ho, W. W. H., Glancy, E. & Eckersley-Maslin, M. A. A low-input high resolution sequential chromatin immunoprecipitation method captures genome-wide dynamics of bivalent chromatin. Epigenetics Chromatin 17, 3 (2024).

53. Vu, H. & Ernst, J. Universal chromatin state annotation of the mouse genome. Genome Biol. 24, 153 (2023).

54. Galindo, G. et al. AI-assisted protein design to rapidly convert antibody sequences to intrabodies targeting diverse peptides and histone modifications. Sci. Adv. 12, eadx8352 (2026).

55. Styles, M. J. et al. PANCS-Binders: a rapid, high-throughput binder discovery platform. Nat. Methods 22, 1720–1730 (2025).

56. Dauparas, J. et al. Robust deep learning–based protein sequence design using ProteinMPNN. Science 378, 49–56 (2022).

57. Watson, J. L. et al. De novo design of protein structure and function with RFdiffusion. Nature 620, 1089–1100 (2023).

58. Pacesa, M. et al. One-shot design of functional protein binders with BindCraft. Nature 646, 483–492 (2025).

59. Stark, H. et al. BoltzGen: Toward Universal Binder Design. Preprint at 10.1101/2025.11.20.689494 (2025).

60. Chai Discovery Team et al. Drug-like antibody design against challenging targets with atomic precision. Preprint at 10.1101/2025.11.29.691346 (2025).

61. Picelli, S. et al. Tn5 transposase and tagmentation procedures for massively scaled sequencing projects. Genome Res. 24, 2033–2040 (2014).

62. Baubec, T., Ivánek, R., Lienert, F. & Schübeler, D. Methylation-Dependent and -Independent Genomic Targeting Principles of the MBD Protein Family. Cell 153, 480– 492 (2013).

63. Grandi, F. C., Modi, H., Kampman, L. & Corces, M. R. Chromatin accessibility profiling by ATAC-seq. Nat. Protoc. 17, 1518–1552 (2022).

64. Bhardwaj, V. et al. snakePipes: facilitating flexible, scalable and integrative epigenomic analysis. Bioinformatics 35, 4757–4759 (2019).

65. Langmead, B. & Salzberg, S. L. Fast gapped-read alignment with Bowtie 2. Nat. Methods 9, 357–359 (2012).

66. Li, H. et al. The Sequence Alignment/Map format and SAMtools. Bioinformatics 25, 2078–2079 (2009).

67. Picard Toolkit. Broad Institute, GitHub repository https://broadinstitute.github.io/picard/ (2019).

68. Ramírez, F. et al. deepTools2: a next generation web server for deep-sequencing data analysis. Nucleic Acids Res. 44, W160–W165 (2016).

69. Team, T. B. D. BSgenome.Mmusculus.UCSC.mm10. Bioconductor 10.18129/B9.BIOC.BSGENOME.MMUSCULUS.UCSC.MM10 (2017).

70. Michael Lawrence, V. C. rtracklayer. Bioconductor 10.18129/B9.BIOC.RTRACKLAYER (2017).

71. P. Aboyoun, H. P. GenomicRanges. Bioconductor 10.18129/B9.BIOC.GENOMICRANGES (2017).

72. Sonali Arora, M. M. GenomeInfoDb. Bioconductor 10.18129/B9.BIOC.GENOMEINFODB (2017).

73. H. Pagès, P. A. Biostrings. Bioconductor 10.18129/B9.BIOC.BIOSTRINGS (2017).

74. Martin Morgan, H. P. Rsamtools. Bioconductor 10.18129/B9.BIOC.RSAMTOOLS (2017).

75. Hervé Pagès, V. O. GenomicAlignments. Bioconductor 10.18129/B9.BIOC.GENOMICALIGNMENTS (2017).

76. Stadler, M. B. et al. DNA-binding factors shape the mouse methylome at distal regulatory regions. Nature 480, 490–495 (2011).

77. Amemiya, H. M., Kundaje, A. & Boyle, A. P. The ENCODE Blacklist: Identification of Problematic Regions of the Genome. Sci. Rep. 9, 9354 (2019).

78. Karimzadeh, M., Ernst, C., Kundaje, A. & Hoffman, M. M. Umap and Bismap: quantifying genome and methylome mappability. Nucleic Acids Res. 10.1093/nar/gky677 (2018) doi:10.1093/nar/gky677.

79. Kagda, M. S. et al. Data navigation on the ENCODE portal. Nat. Commun. 16, 9592 (2025).

