## Supplementary material for "Read&Tag enables antibody-independent combinatorial and multifactorial chromatin profiling": Schneider_et_al_supplementary_figures_biorxiv_2026

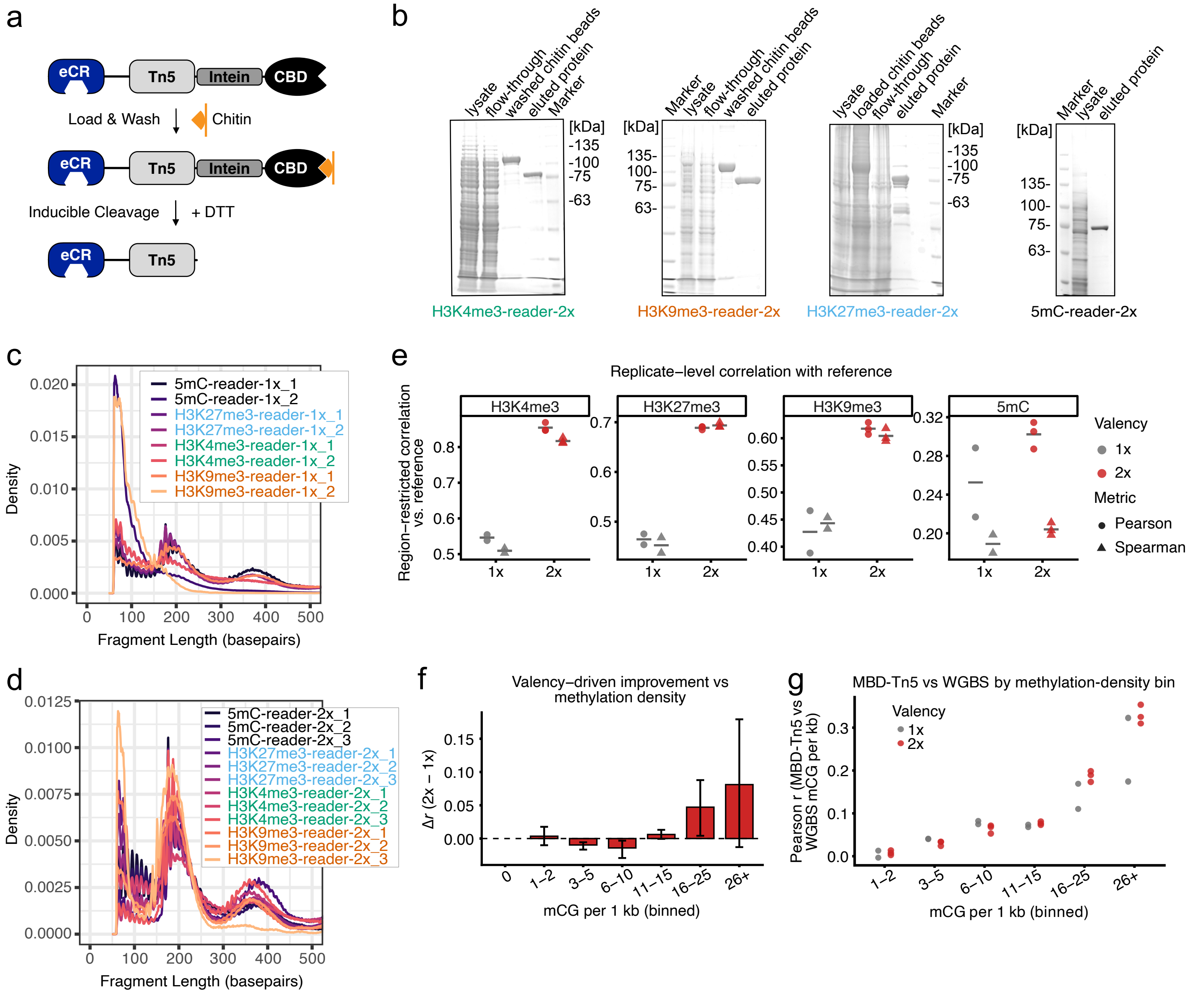

**Supplementary Figure 1. Design, purification, and basic characterization of eCR-Tn5 fusion proteins.**
**(a)** Overview of the intein-mediated purification strategy. eCR-Tn5 fusion proteins are expressed in bacteria, immobilized on chitin resin via the CBD, and released by inducible intein cleavage upon addition of dithiothreitol (DTT), yielding tag-free eCR-Tn5 proteins. **(b)** Coomassie-stained SDS-PAGE gels showing representative purification steps for different eCR-Tn5 constructs, including TAF3-PHD-x2-Tn5 (H3K4me3-reader-2x), CBX1-Chromo-x2-Tn5 (H3K9me3-reader-2x), CBX2-ChromoVD-x2-Tn5 (H3K27me3-reader-2x), and MBD1-MBD-x2-Tn5 (5mC-reader-2x). Lanes show bacterial lysate, flow-through, washed chitin beads, and eluted protein fractions, with molecular weight markers indicated. **(c, d)** Fragment length distributions of sequencing libraries generated using different eCR-Tn5 fusion proteins. Density plots display aggregate fragment length profiles, illustrating the size ranges obtained from reader-guided tagmentation. **(e)** Region-restricted correlation between each Read&Tag replicate and its reference, for single-domain (1x; grey; n=2) and dual-domain (2x; red; n=3) constructs across four marks. Pearson r (circles) and Spearman ρ (triangles) are shown; grey bars are the replicate mean per valency and metric. Each panel has an independent y-axis, so values are not comparable across marks. Correlations use 1-kb windows overlapping each mark's reference region set (24,255-846,071 windows). For histone marks, MACS2 replicate-consensus peaks correlated against matched ChIP-seq. For 5mC, top-tertile WGBS methylated-CpG-density windows (excluding CpG islands) correlated against WGBS density. For every histone mark, all 2x replicates exceed all 1x replicates in both metrics. **(f, g)** Depth-matched density-stratified analysis of MBD-Tn5 versus WGBS mCG density. All five MBD libraries were subsampled to a common usable-fragment depth. QC-passed non-CpG-island windows (n = 2,196,516) were grouped into seven density bins. **(f)** Per-bin Δr (2x − 1x) from all 2 × 3 = 6 cross-replicate pairings, as mean (bar) and [min, max] range (whiskers); the pairings are non-independent, so whiskers are descriptive, not a confidence interval. The mCG = 0 bin has undefined correlation (zero variance), shown for completeness. The 2x advantage is reproducible at high density (16-25 mCpG/kb). **(g)** Underlying per-replicate Pearson r (1x grey, 2x red) for the six defined bins.

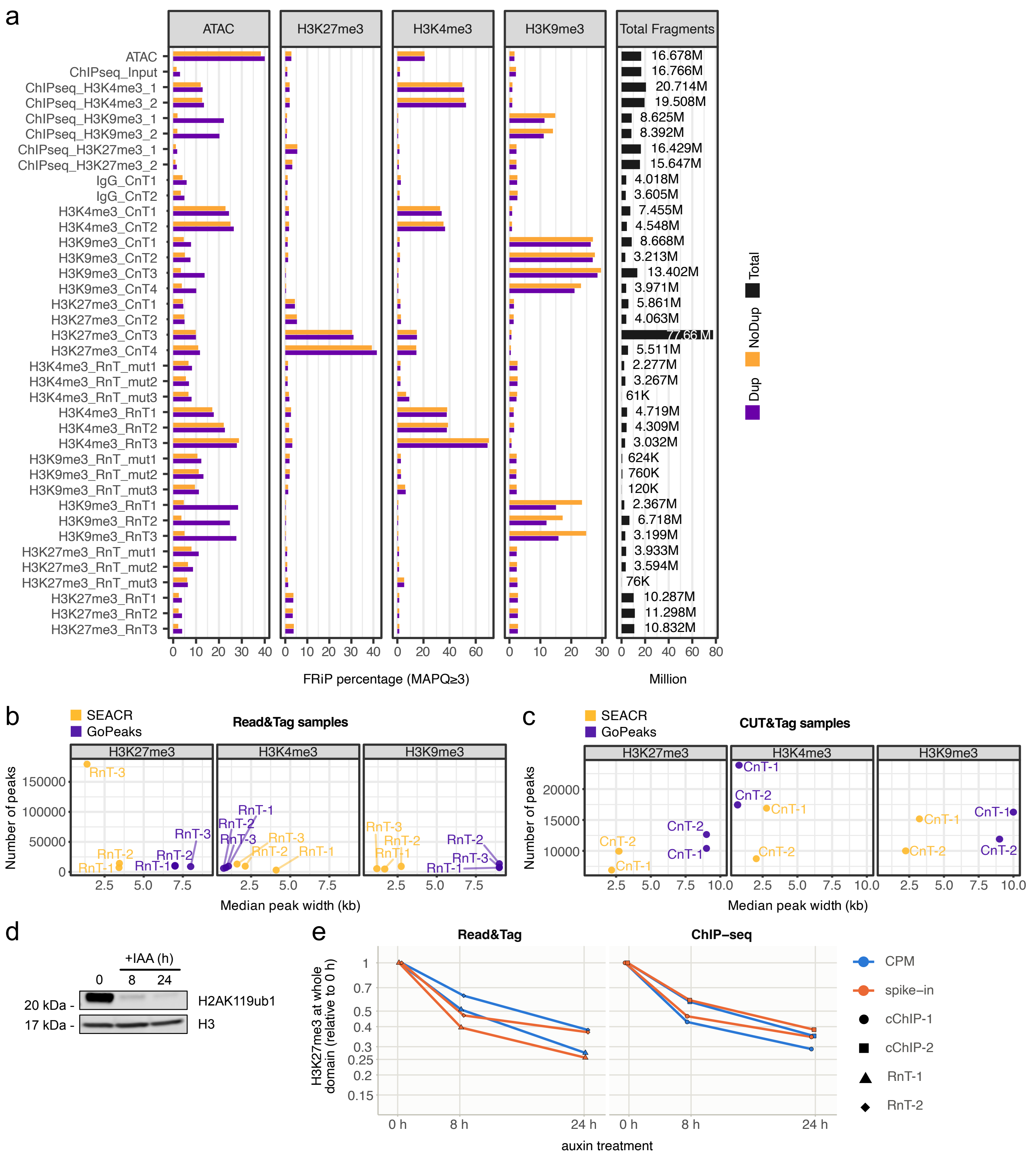

**Supplementary Figure 2. Quality metrics and peak characteristics of Read&Tag datasets across three histone modifications.**
Summary of sequencing-based quality metrics and peak properties for Read&Tag datasets targeting H3K27me3, H3K4me3, and H3K9me3, alongside reference datasets. **(a)** Bar plots showing the fraction of paired reads (fragments) in peaks (FRiP) for individual datasets across chromatin assays. Panels are organized by assay type (ATAC, H3K27me3, H3K4me3, and H3K9me3), with each row corresponding to a dataset, including Read&Tag (RnT), CUT&Tag (CnT, refs. 7, 32-34), ChIP-seq, input, and mutant-reader controls. FRiP values are calculated relative to reference peak sets. Duplicated (Dup) and non-duplicated (No Dup) fragments with a mapping quality score of 3 or higher (MAPQ≥3) are displayed separately. A column on the right indicates total fragments (MAPQ≥3) for each dataset. **(b-c)** Scatter plots summarizing peak characteristics for Read&Tag (b) and CUT&Tag (c; ref. 33) datasets across histone modifications. Each point represents an individual dataset, with the x-axis indicating median peak width (kb) and the y-axis indicating the total number of peaks identified. Separate panels are shown for H3K27me3, H3K4me3, and H3K9me3, and points are labeled by biological replicate and peak-calling strategy (GoPeaks or SEACR). **(d)** Western blot analysis showing reduction of H2AK119ub1 in the PRC1-degron cell line. Histone H3 serves as loading control. Identical results were obtained from two independent experiments. **(e)** H3K27me3 loss following PRC1 degradation, measured by Read&Tag (left) and by spike-in-calibrated ChIP-seq (right; Dobrinić et al., reprocessed through an identical pipeline). Degron mESCs were treated with auxin for 0, 8 and 24 hours. For each replicate and normalization, signal was quantified over 2,745 H3K27me3 whole-domains called from the untreated reference ChIP-seq independently of Read&Tag and is plotted as the median of per-domain ratios to the matched 0 h sample (log scale). Symbols denote biological replicates (RnT-1, RnT-2; cChIP-1, cChIP-2), so the separation between paired symbols is the replicate spread; colour denotes normalization, CPM (blue) or Drosophila melanogaster spike-in (orange).

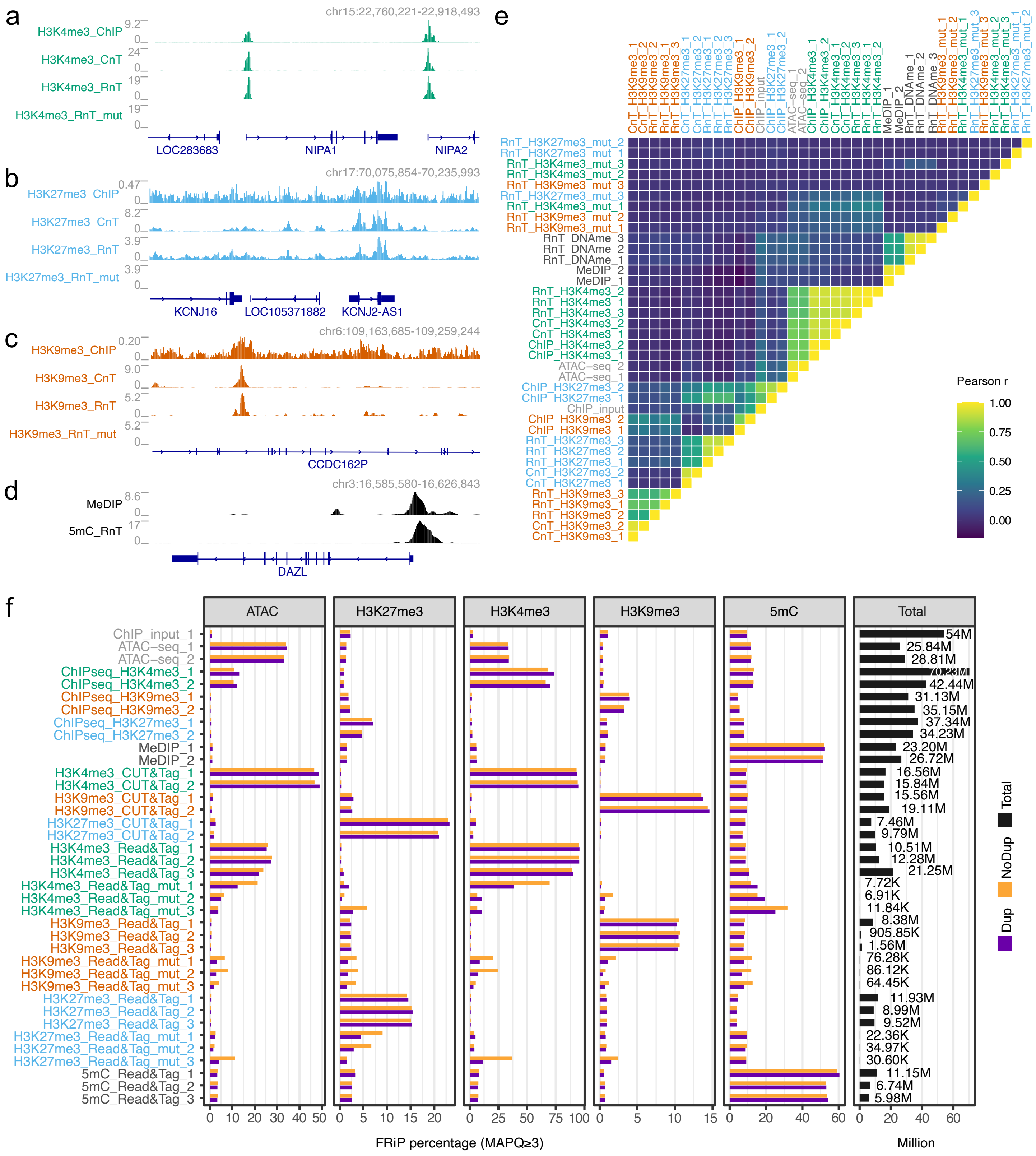

**Supplementary Figure 3. Read&Tag faithfully captures chromatin modification patterns in human MCF7 cells.** Read&Tag profiling of three histone modifications and 5mC in human MCF7 cells and comparison to published ChIP-seq and CUT&Tag datasets using locus-level, genome-wide, and peak-based metrics.

**(a-d)** Genome browser tracks at representative loci showing average profiles for H3K4me3 (a), H3K27me3 (b), H3K9me3 (c), and 5mC (d), obtained from 250,000 MCF7 cells using Read&Tag. Corresponding reference ChIP-seq (ref. 39), CUT&Tag (ref. 40), and MeDIP (ref. 41) tracks are shown for comparison, with gene annotations displayed below regions.

**(e)** Pairwise Pearson correlation matrix comparing genome-wide signal distributions across Read&Tag biological replicates (n=3) and reference datasets (n=2) for the three histone modifications and 5mC. Pearson correlations are shown for individual reader-Tn5 configurations and replicates.

**(f)** Fraction of paired reads (fragments) in peaks (FRiP) for Read&Tag datasets targeting H3K27me3, H3K4me3, H3K9me3, and 5mC (DNAme). Bar plots show the percentage of sequencing fragments with a mapping quality score of 3 or higher (MAPQ≥3) overlapping ATAC-, MeDIP-, or ChIPseq-defined reference peak sets, with duplicated (Dup) and non-duplicated (No Dup) fragments indicated separately. A column on the right indicates total fragments (MAPQ≥3) for each dataset.

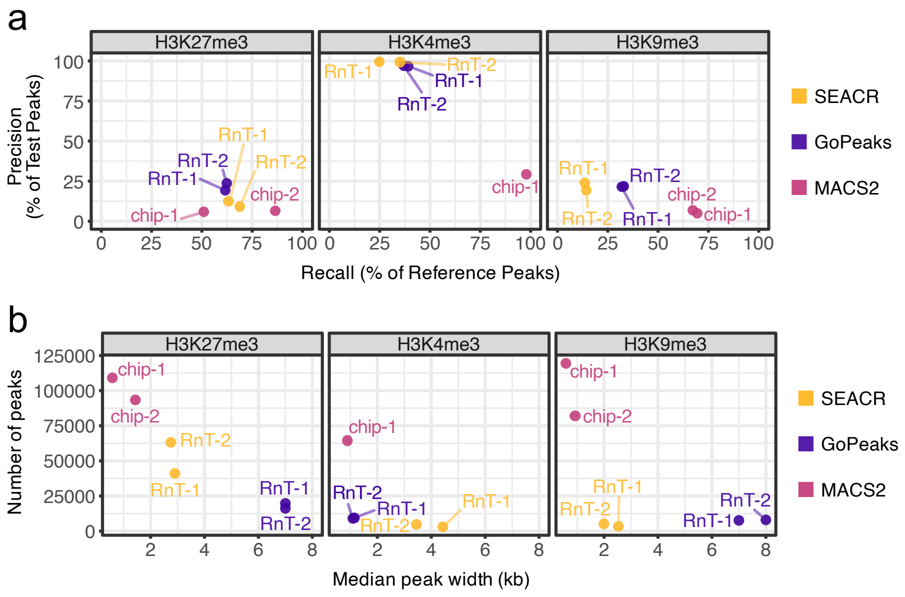


**Supplementary Figure 4. Comparison of peak-calling strategies applied to Read&Tag datasets in liver tissue.**
**(a)** Precision-recall plots summarizing peak recovery for Read&Tag datasets across the three ENCODE reference peak sets. Precision and recall values are computed relative to ENCODE reference peak sets, with points labeled by ChIP-seq (refs. 43, 44), and Read&Tag configuration, biological replicate, and peak-calling strategy (GoPeaks, SEACR, or MACS2). Precision is defined as the fraction of called peaks overlapping reference peaks and recall defined as the fraction of reference peaks recovered.
**(b)** Scatter plots summarizing peak characteristics for each histone modification and peak-calling strategy. Each point represents an individual dataset and is positioned according to the number of identified peaks (y-axis) and the median peak width in kilobases (x-axis). Separate panels are shown for H3K27me3, H3K4me3, and H3K9me3, with peak-calling method indicated by color.

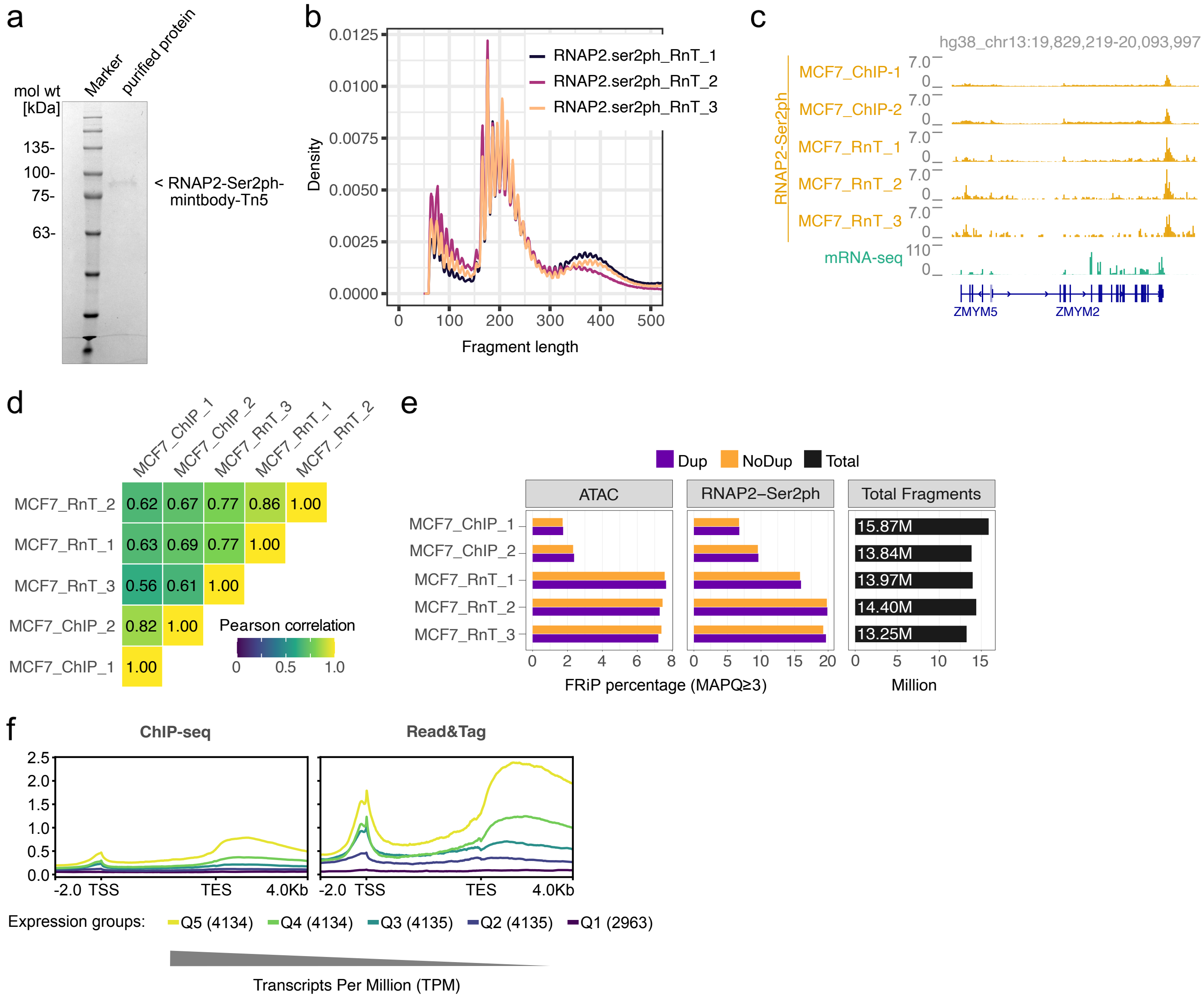


**Supplementary Figure 5. Production, library characteristics, and performance of RNAP2-Ser2ph Read&Tag in human MCF7 cells.**

**(a)** Coomassie-stained SDS-PAGE gel showing the purified RNAP2-Ser2ph reader-Tn5 fusion protein. A molecular weight marker is shown alongside the purified protein, with the expected size of the fusion indicated.

**(b)** Fragment length distributions of sequencing libraries generated by RNAP2-Ser2ph profiling using Read&Tag in mouse ES cells. Density plot displays aggregate fragment length profiles across biological replicates, illustrating the range and distribution of fragment sizes obtained.

**(c)** Genome browser tracks at representative loci in the human genome (hg38) showing RNAP2-Ser2ph signal obtained by ChIP-seq (ref. 48), and Read&Tag in MCF7 cells, displayed together with mean mRNA-seq signal (n=3).

**(d)** Pairwise Pearson correlation matrix comparing genome-wide RNAP2-Ser2ph signal distributions obtained by ChIP-seq (ref. 48), and Read&Tag across biological replicates.

**(e)** Bar plots showing the fraction of duplicated (Dup) and non-duplicated (No Dup) fragments with a mapping quality score of 3 or higher (MAPQ≥3) overlapping ChIP-seq-defined peaksets (MACS2) for RNAP2-Ser2ph and accessibility-associated regions generated by ATAC-seq (ref. 39) in MCF7 cells. A column on the right indicates total fragments (MAPQ≥3) for each dataset.

**(f)** Metagene profiles of RNAP2-Ser2ph mean signals across the bodies of protein-coding genes (hg38), aligned at transcription start sites (TSS) and transcription end sites (TES). Genes are grouped by mRNA-seq expression level, with Q1 representing transcriptionally silent genes (TPM = 0) and Q2-Q5 representing equal-sized quantiles of expressed genes (TPM > 0). Profiles are shown for published ChIP-seq (n=2; ref. 48), and Read&Tag (n=3) mean signals to illustrate transcription-associated signal patterns across expression strata.

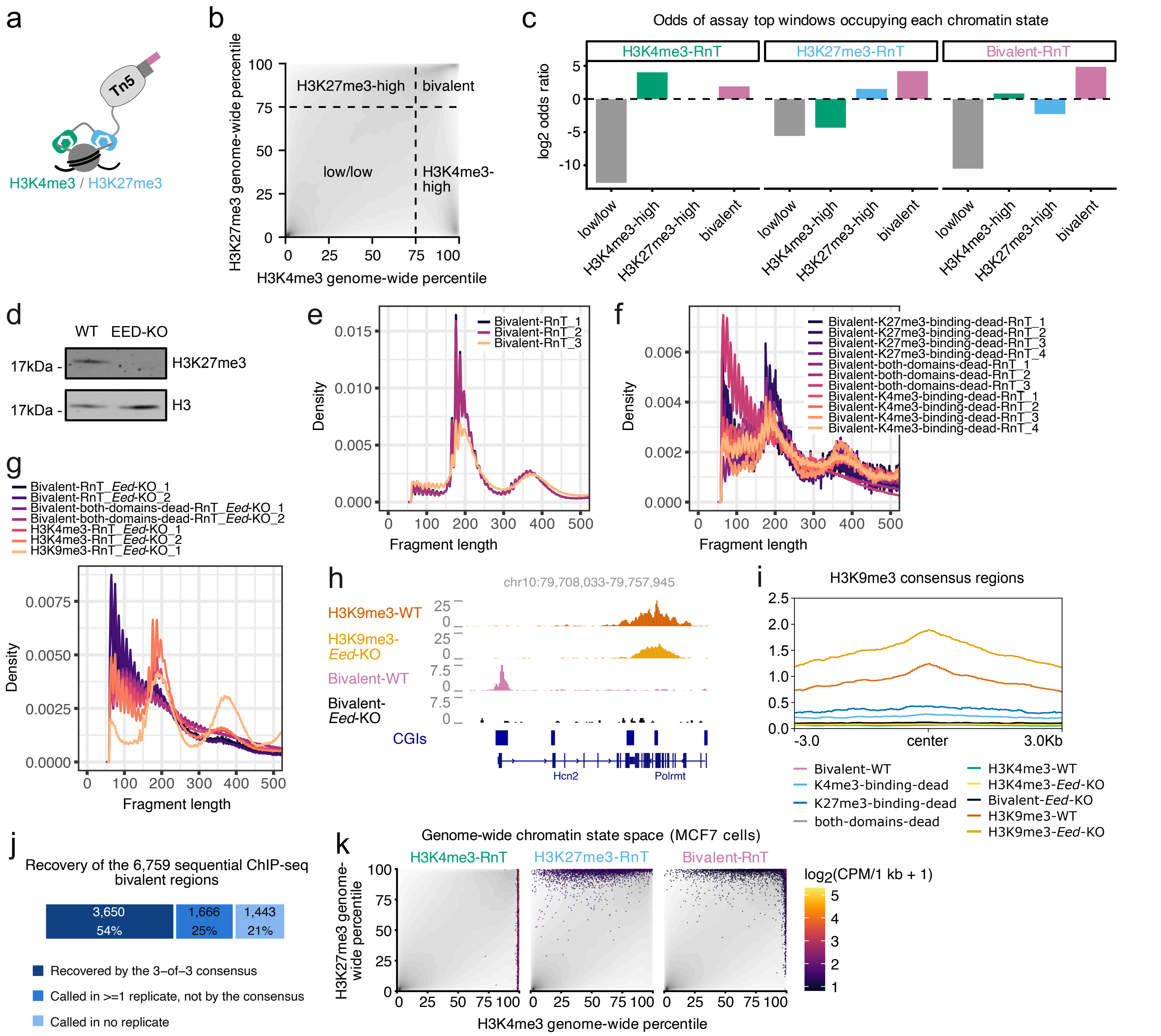


**Supplementary Figure 6. Characterization and benchmarking of combinatorial Read&Tag datasets across chromatin states.**

**(a)** Schematic illustration of the combinatorial reader-Tn5 fusion used to target nucleosomes carrying both H3K4me3 and H3K27me3 modifications.

**(b)** Schematic representation of chromatin-state space defined by percentile-ranked H3K4me3 (x-axis) and H3K27me3 (y-axis) signals. The plot illustrates four discrete chromatin-state categories (low/low, H3K4me3-high, H3K27me3-high, and bivalent) used for downstream classification and analysis.

**(c)** Bar plots summarizing the distribution of top-ranked assay windows across chromatin-state categories for H3K4me3-, H3K27me3-, and bivalent Read&Tag datasets. Values are expressed as log2 odds ratios derived from contingency-based comparisons between assay-enriched windows and genome-wide background distributions.

**(d)** Western blot analysis showing loss of H3K27me3 in the *Eed*-KO cell line. Histone H3 serve as loading control. Identical results were obtained from two independent experiments.

**(e-f)** Fragment length distributions of sequencing libraries generated by Read&Tag using the combinatorial reader-Tn5 fusion in wildtype cells (E), the “both-dead-” and two “single-dead-domain” combinatorial chromatin reader-Tn5 constructs wildtype cells (F), or the combinatorial-, the H3K4me3-, and H3K9me3-reader-Tn5 fusions and in H3K27me3-deficient cells (G). Density plots display aggregate fragment length profiles across biological replicates, illustrating the range and distribution of fragment sizes obtained.

**(h)** Genome browser view at a representative locus showing Read&Tag profiles in wildtype (WT) versus H3K27me3-deficient cells (*Eed*-KO). Tracks include average signals for H3K9me3 Read&Tag profiles in WT cells (n=3), H3K9me3 Read&Tag profile in *Eed*-KO cells (n=1), combinatorial Read&Tag profiles in WT cells (n=3), and combinatorial Read&Tag profiles in *Eed*-KO cells (n=2). Gene models and CpG islands (CGIs) are indicated below the tracks.

**(i)** Aggregate signal profiles centered on consensus peak regions for H3K9me3 Read&Tag in wildtype cells. Plot shows average signal intensities across genomic windows spanning ±3 kb around peak centers. Separate curves are shown for each modality.

**(j)** Stacked bar partitioning the 6,759 consensus bivalent regions defined by sequential ChIP-seq according to how combinatorial Read&Tag detected them: recovered by the 3-of-3 replicate consensus, called in at least one replicate but not by the consensus, or called in no replicate. Overlap was scored as ≥1 shared base pair.

**(k)** Genome-wide projection of 1 kb genomic windows into chromatin-state space defined by percentile-ranked H3K4me3 (x-axis) and H3K27me3 (y-axis) ChIP-seq signals in MCF7 cells. The background grayscale density represents the distribution of all genomic windows, while colored points denote the top 1% highest-signal windows for each Read&Tag assay. Color intensity reflects Read&Tag signal (log2 CPM per 1 kb + 1).

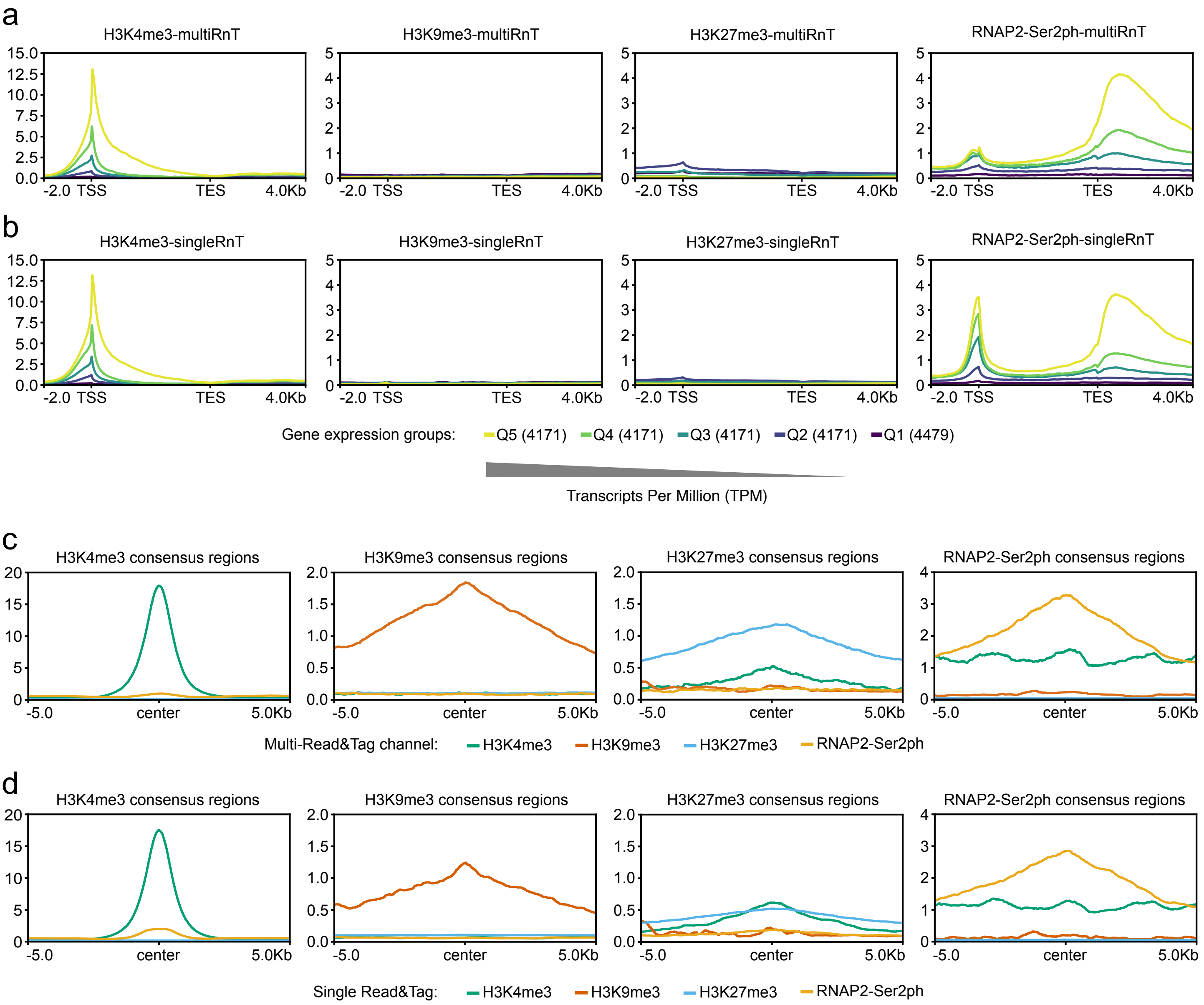


**Supplementary Figure 7. Within-target response and target specificity of multiplexed Read&Tag.**
**(a,b)** Metagene profiles displaying average Multi-Read&Tag (a) or single-target Read&Tag (b; singleRnT) signals across the bodies of protein-coding genes (mm10) for H3K4me3, H3K9me3, H3K27me3, and RNAP2-Ser2ph. Signals are aligned relative to transcription start sites (TSS) and transcription end sites (TES), with genomic coordinates scaled to include upstream and downstream flanking regions. Genes are grouped by mRNA-seq expression level, with Q1 representing transcriptionally silent genes (TPM = 0) and Q2-Q5 representing equal-sized quantiles of expressed genes (TPM > 0). The average signals of two Multi-Read&Tag and three single-target Read&Tag biological replicates are shown in the profiles.
**(c,d)** Aggregate signal profiles centered on consensus peak regions for H3K4me3, H3K9me3, H3K27me3, and RNAP2-Ser2ph in Multi-Read&Tag (c) and single-target Read&Tag (d). Plots show average signal intensity across genomic windows spanning ±5 kb around peak centers. Separate curves are shown for individual Read&Tag datasets corresponding to H3K4me3, H3K9me3, H3K27me3, and RNAP2-Ser2ph, allowing comparison of signal distributions across marks within the same genomic regions. Consensus peak sets (GoPeaks) were defined from single-target Read&Tag replicates (n=3) alone and are therefore independent of the multiplexed data.
